# Locus coeruleus-related insula activation is associated with implicit learning

**DOI:** 10.1101/2025.05.10.653174

**Authors:** Martin J. Dahl, Tiantian Li, Mara Mather, Markus Werkle-Bergner

**Author notes:** **Corresponding author**: MJD. These authors contributed equally (shared senior authorship) and are listed alphabetically (Mara Mather, Markus Werkle-Bergner).

## Abstract

The noradrenergic locus coeruleus and its neuromodulatory cortical projections are critical for adaptive behavior, however how they contribute to human implicit learning in novel environments is not yet well understood, due to the challenges of assessing this system non-invasively. Here, we combined multimodal neuroimaging—including locus-coeruleus-sensitive structural MRI, concurrent pupillometry–EEG, pupillometry–fMRI, and PET-derived noradrenergic transporter maps—with repeated behavioral assessments to investigate noradrenergic contributions to implicit learning across younger and older adults (n = 77). Salient expectation-violating stimuli elicited pupil dilation, indicating enhanced neuromodulation, activated the action-mode network and deactivated the default-mode network. Pupil-linked BOLD responses suggested a functional coupling between the locus coeruleus and action-mode network, further supported by spatial overlap of activation patterns with PET-derived noradrenergic transporter maps. Locus coeruleus MRI-guided functional connectivity analyses demonstrated coupled locus coeruleus and anterior insula activity, suggesting a noradrenergic role in shifting cortical dynamics toward action-oriented processing. Behaviorally, participants implicitly learned the statistical task structure over time, shifting their response times based on stimulus probabilities. Critically, stronger locus coeruleus integrity, greater task-related anterior insula activation, and more pronounced pupil dilation were associated with enhanced implicit learning, highlighting the behavioral relevance of noradrenergic neuromodulation. Notably, noradrenergic responses and their link to learning were preserved across age groups, suggesting a robust noradrenergic role in supporting adaptive behavior throughout adulthood. Complementary electrophysiological analyses revealed pupil-linked increases in cortical excitability that were coupled with shifts from default-mode to action-mode network activation. Together, these findings provide novel insights into the neuromodulatory mechanisms underlying learning and cognitive flexibility, emphasizing the pivotal role of locus coeruleus–action-mode network interactions in behavioral adaptation.

## 1. Introduction

Optimal behavior requires dynamic adaptation to changing environments^1^. On a neurochemical level, unexpected environmental changes transiently increase noradrenaline release from the locus coeruleus, a brainstem nucleus that facilitates attention and learning^2–4^. Specifically, noradrenaline release has long been conceptualized as a neural interrupt signal that promotes rapid behavioral adaptation to new environmental imperatives^5–8^. The momentary processing of salient stimuli is thereby facilitated by a noradrenergic enhancement of neural gain, effectively increasing signal-to-noise ratios in neural circuits^2,9–12^. By contrast, long-term adjustments to salient events are mediated by noradrenergic influences on hippocampal synaptic plasticity^3,13–16^.

In a similar vein, more recent accounts propose that the locus coeruleus is activated when predictions about the world are violated. The resulting noradrenaline release helps refine internal models of the world by adjusting the rate of learning in cortical target regions^17–21^, but direct evidence from humans is scarce.

To enable these functions, the locus coeruleus receives critical contextual information about stimulus salience and utility from cortical structures such as the anterior cingulate and insula^2,9^. In primates, these forebrain structures send major direct inputs to the locus coeruleus to recruit noradrenergic neuromodulation and modulate brain-wide processing according to environmental demands^9^.

The anterior cingulate and insular cortices have recently been grouped into the action-mode network^22^, overlapping with the cingulo-opercular network or ventral attention network^6,23,24^. The action-mode network is closely linked to initiating and maintaining states of heightened arousal and focused attention, as well as the generation and updating of actions based on salient internal or external signals^22,25^. In this context, arousal is defined as a continuum of sensitivity to environmental stimuli^26^, tightly regulated by noradrenergic neuromodulation^27–29^. In line with this, it has been proposed that the locus coeruleus modulates or is part of the action-mode network^6,22,30^, but direct in vivo evidence remains limited.

Aging dysregulates the noradrenergic system^31–33^, possibly due to the age-related accumulation of pathology^34–37^. In accordance with this, converging evidence based on cognitive assessments suggests that age differences in proxies of noradrenergic neuromodulation contribute to late-life impairments in attention^38–40^, learning, and memory^41–43^; i.e., in situations when participants were *explicitly* instructed about the cognitive testing. However, the role of the locus coeruleus in how younger and older adults *implicitly* learn to adapt their behavior in new situations remains underexplored.

Here, we investigated the implications of locus coeruleus–cortical network interactions for implicit behavioral adaptation using a multimodal age-comparative approach. To this end, we used a modified conditioned oddball task, previously shown to reliably increase locus coeruleus spiking in non-human primates^44^. In this variant of the oddball task, participants detect infrequent, distinctive stimuli amid a series of repetitive standard stimuli. To further increase recruitment of the locus coeruleus, infrequent oddball stimuli are pre-conditioned by repeatedly pairing them with appetitive or aversive outcomes^40,44–46^. Our participants completed the conditioned oddball task twice on consecutive days, responding with different button presses depending on stimulus identity (see Figure 1). Trial-level behavioral responses from the task allowed us to examine implicit learning within and across sessions. In this context, implicit learning refers to how initially naïve participants pick up the statistical regularities of the originally new but increasingly familiar task environment and optimize their behavior over time (i.e., without explicit instructions).

**Figure 1.**
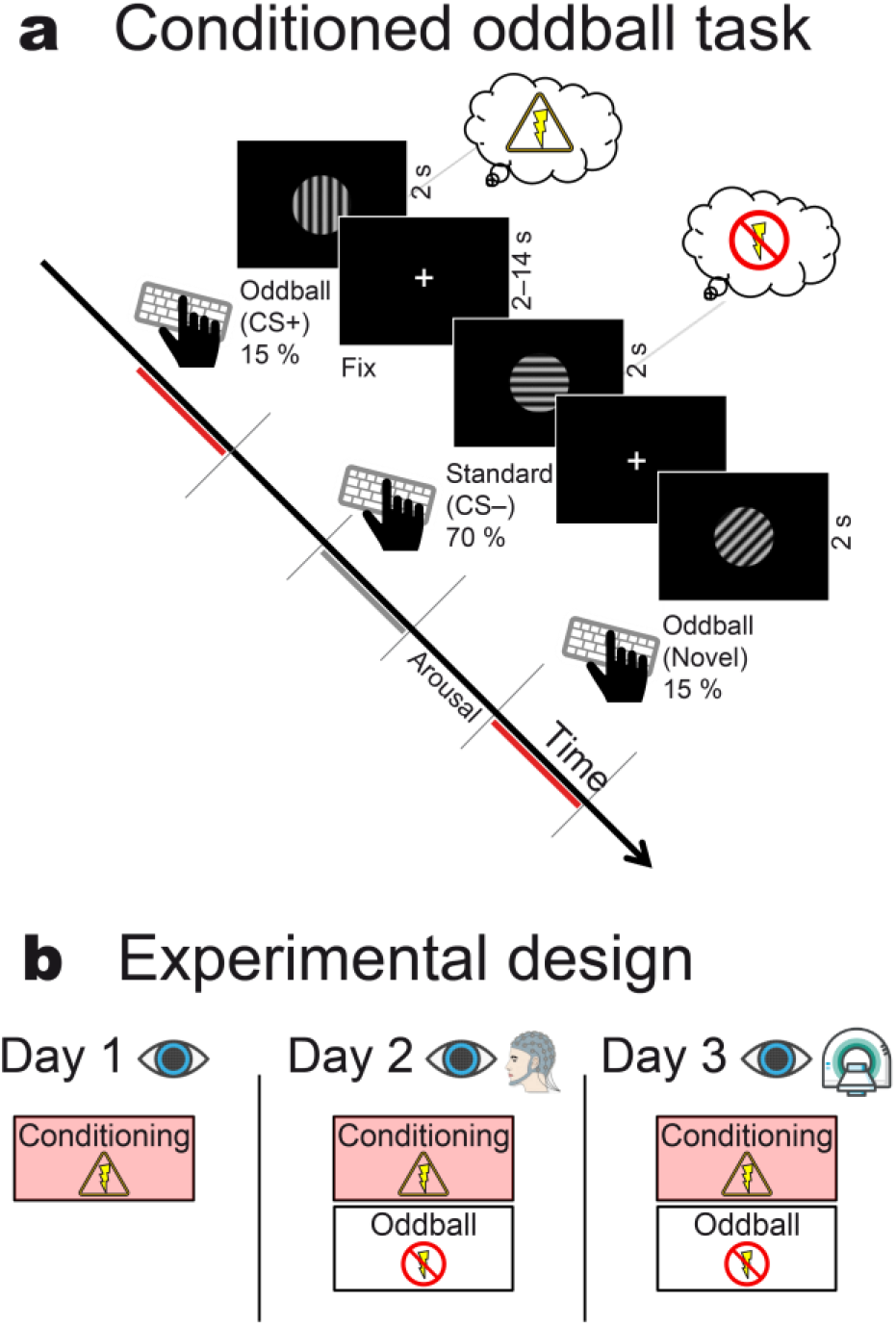
Overview of the conditioned oddball task (a) and experimental phases (b). As part of a three-stimulus oddball task, previously conditioned stimuli (CS+) were presented as infrequent oddball stimuli while non-conditioned perceptually matched control stimuli (CS–) served as frequent standards. A second oddball category consisted of perceptually matched but previously not shown stimuli, hypothesized to elicit a novelty response. During the oddball tasks, participants were instructed to respond as accurately and quickly as possible with a different button press based on the orientation of the shown stimulus. (**b**) Fear conditioning sessions were repeated over three consecutive days while collecting eye tracking and EEG data. The conditioned oddball task took place on Day 2 (pupillometry-EEG) and Day 3 (pupillometry-fMRI). During the oddball tasks, no electrical stimulation was applied.

To overcome challenges in non-invasive assessment posed by the locus coeruleus’ small size and location in the brainstem^47–49^, we relied on several recent additions to the neuroscience toolkit. In particular, dedicated structural magnetic resonance imaging (MRI) sequences can reveal the locus coeruleus as a cluster of distinct bright (hyperintense) voxels bordering the fourth ventricle^41,49^. Transgenic animal^50,51^ and human post-mortem^52^ validation studies suggest this signal may serve as an in-vivo proxy for the integrity of the locus and other neuromodulatory nuclei. Here, we relied on structural neuromodulation-sensitive MRI to assess individual differences in locus coeruleus integrity and to reliably localize the small brainstem nucleus in functional analyses. To additionally track locus coeruleus activation patterns, we combined fMRI with concurrent pupillometry, as increasing arousal-related neuromodulatory activity, including in the noradrenergic system, dilates pupils^53–55^. Finally, we compared the observed pupillometry–fMRI activation maps to the brain-wide distribution of noradrenergic and other neuromodulatory transporters^56^ to gauge potential dependencies. Complementary electrophysiological analyses focused on the P300, a positive parietal event-related potential, that is influenced by locus coeruleus activity^57,58^, as well as markers for cortical excitability that are sensitive to neuromodulatory activation^4,39,59– 61^. In sum, younger and older adults completed a conditioned oddball task while we assessed multiple proxies for locus coeruleus structure and function to explore their role in implicit learning.

In a brief synopsis of our findings, salient stimuli reliably dilated younger and older participants’ pupils and activated the action-mode network. Pupil-indexed neuromodulation correlated with brainstem and action-mode network activation, and was associated with noradrenergic transporter distribution, suggesting potential links to the locus coeruleus, which we confirmed using functional connectivity analyses. Initially naïve participants implicitly learned the structure of the new task environment and adapted their behavior over time. Behavioral adaptation (i.e., implicit learning) was positively associated with structural and functional locus coeruleus and action-mode network measures. Complementary electrophysiological analyses revealed pupil-linked increases in cortical excitability that were coupled to large-scale fMRI network transitions, including action-mode network activation. Together, these results suggest that noradrenergic signals support refining internal models of environmental contingencies for behavioral adaptation by modulating attention and memory following salient events. By integrating advanced MRI, pupillometry, EEG, and functional connectivity analyses, we reveal critical insights into how neuromodulatory signals dynamically drive behavioral flexibility.

## 2. Results

### Conditioning reliably increases pupil-indexed neuromodulation across age groups

To increase recruitment of the locus coeruleus, experimental stimuli that were subsequently shown in the main task were pre-conditioned by repeatedly pairing them with an aversive electrical stimulation (Figure 1,^44^). Robust pupil dilation following presentation of conditioned stimuli (CS+) relative to perceptually matched control stimuli (CS–) suggests this procedure enhanced neuromodulatory activation, including in the noradrenergic system^55^, across age groups and assessment days (Figure 2a; all *p*_cluster-corr_ ≤ 0.002).

**Figure 2.**
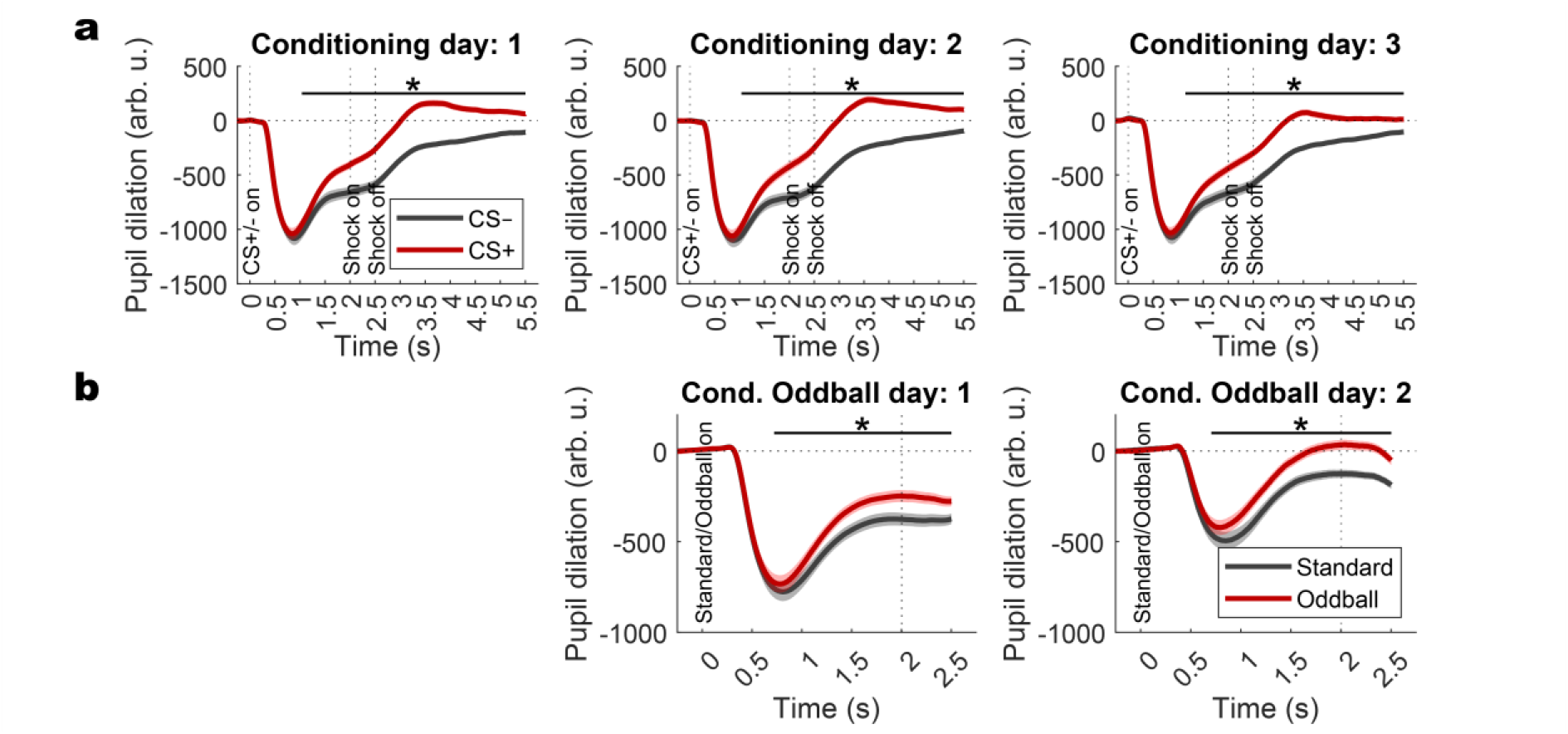
Stimulus-related pupil responses during fear conditioning and conditioned oddball tasks. (**a**) During daily fear conditioning sessions, fear conditioned stimuli (CS+; red) elicited stronger pupil dilation relative to perceptually matched control stimuli (CS–, black), suggesting neuromodulatory activation^53,54^. Note that both stimuli initially elicited comparable pupil constriction due to higher luminance compared to the background (pupil light reflex after stimulus presentation at time 0). (**b**) During subsequent pupillometry-EEG (day 1) and pupillometry-fMRI (day 2) conditioned oddball tasks, oddball presentation (red) was linked to larger pupil dilation relative to perceptually matched control stimuli (standard; black). Differences in pupil light reflexes between the two acquisition days resulted from different background luminance in the two settings (EEG, MRI). The black horizontal lines indicate the temporal extent of the significant group level cluster (*), controlling for multiple comparisons, which is comparable across days. Shaded areas represent standard errors of the mean. Pupil size is expressed in arbitrary units (arb. u.), relative to a pre-stimulus baseline period.

### Conditioned oddball stimuli dilate the pupil and activate the action-mode network

During the subsequent main task, salient stimuli—oddballs that were either pre-conditioned or novel—reliably dilated participants’ pupils (*p*_cluster-corr_ ≤ 0.002) and activated the action-mode network relative to perceptually matched control stimuli (standards [CS–]; Figures 2b and 3).

**Figure 3.**
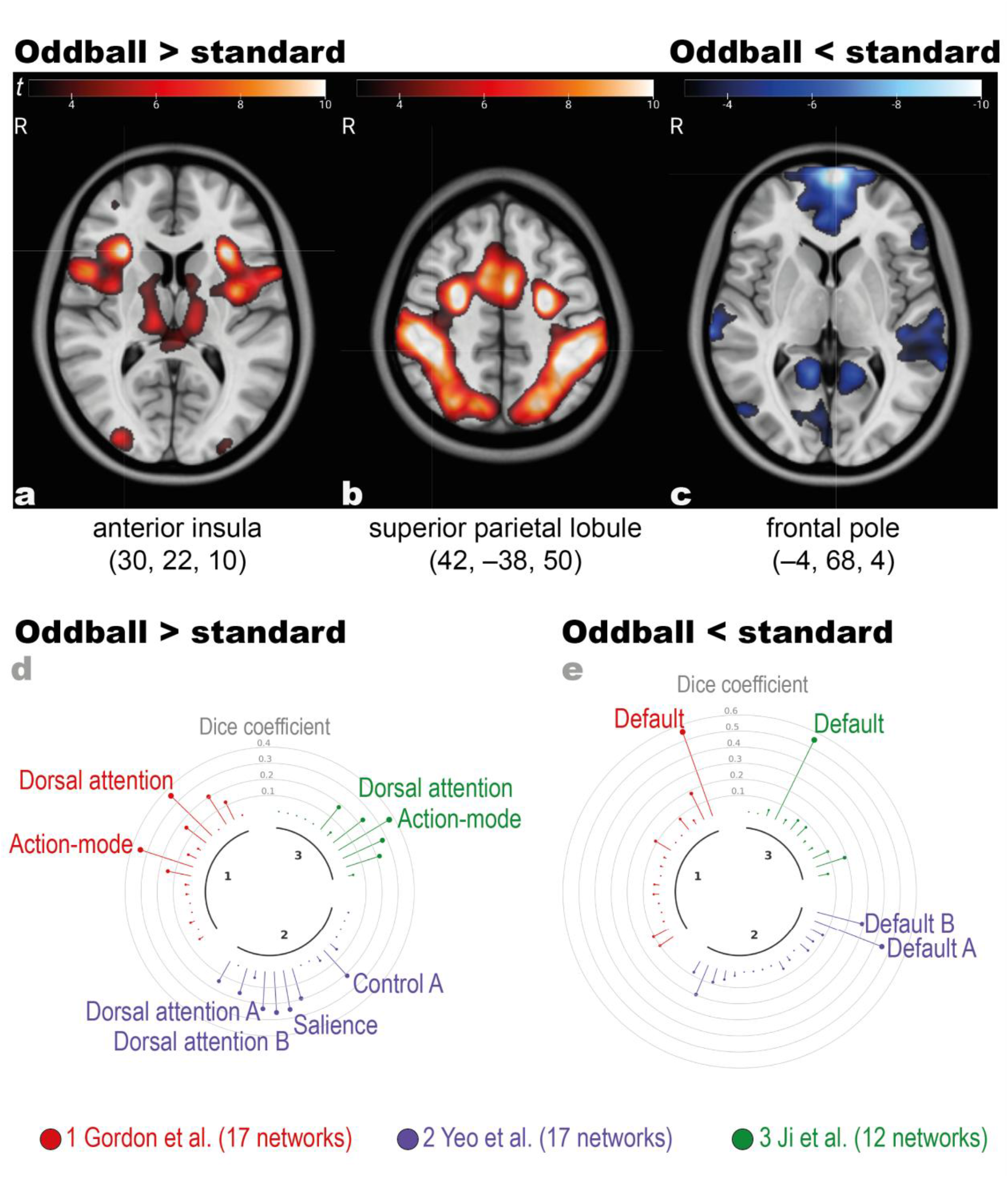
Salient oddball stimuli elicit action-mode network (a) and dorsal attention network activation (b) and deactivate the default mode network (c). For visualization, group contrast maps (oddballs vs standards) are thresholded at *t =* 3, main text results report family-wise error corrected statistics. Contrast maps are available at: osf.io/4t5hj. Numbers below the labeled regions indicate MNI XYZ-coordinates. **(d-e)** Overlap of group-level activation **(d)** and deactivation **(e)** maps with three published functional network atlases^23,63,64^, quantified as Dice coefficients. Only networks with a significant overlap based on permutation tests^24^ and Dice coefficients > 0.2 are labeled. Note that the *Action-mode network* has previously been termed *Cingulo-opercular network*^22^ and the *Salience network* has also been termed *Ventral attention network*^24^.

Action-mode network recruitment encompassed the bilateral anterior insula (*t =* 11.81, *p*_FWE-corr_ < 0.001 MNI: 30, 22, 10), middle cingulate gyrus (*t =* 8.95, *p*_FWE_ < 0.001 MNI: 3.5, 7.5, 31.5), thalamus (*t =* 7, *p*_FWE_ < 0.001 MNI: 12, –20, 12) and cerebellum (*t =* 7.67, *p*_FWE_ < 0.001 MNI: 30, –50, –24;^22,25^). In addition, salient stimuli elicited prominent dorsal attention-network activation (bilateral superior parietal lobule; *t =* 13.8, *p*_FWE-corr_ < 0.001 MNI: 42, –38, 50;^6,25^) and default-mode network *de*activation (e.g., bilateral frontal pole, *t =* –11.64, *p*_FWE_ < 0.001 MNI: –4, 68, 4;^22^). The un/thresholded whole-brain contrast maps are available at osf.io/4t5hj.

Together, this suggests task-related neuromodulatory activation, as indicated by pupil dilation, as well as activation of the action-mode network, reflecting major cortical in- and output regions of the noradrenergic system^6,9,62^. This may indicate that in response to salient expectation-violating stimuli, there is a transition from default-mode to action-mode network activation, likely facilitated by noradrenergic neuromodulation, which supports coping with changing task demands^5,17,22^.

### Pupil-indexed neuromodulation is associated with action-mode network and brainstem activation

By integrating pupil time series into the fMRI analyses, we directly probe the relationship between the detected cortical activation patterns and markers of neuromodulation. This approach allowed us to test which brain regions’ activity correlated with moment-to-moment pupil fluctuations, which are likely driven by release of neuromodulatory neurotransmitters^53–55^.

Supporting an association between noradrenergic neuromodulation and action-mode activation, network hubs such as bilateral anterior insula (*t =* 7.31, *p*_FWE-corr_ < 0.001 MNI: – 32, 20, –8), middle cingulate cortex (*t =* 5.55, *p*_FWE-corr_ < 0.001 MNI: –4 –0 44), putamen (*t* = 8.52, *p_FWE-corr_* < 0.001 MNI: –18.5, 13.5, –6.5), and thalamus (*t =* 7.88, *p*_FWE-corr_ < 0.001 MNI: 14 –12 10) scaled their activity with pupil size (Figure 4). The un/thresholded whole-brain contrast maps are available at osf.io/4t5hj. Supplementary Figure S1 extends this analysis to include variable delays between BOLD and pupil data and Supplementary Figure S2 shows the auto-correlation of the two signals. These findings corroborate recent observations—linking human intracranial insula recordings with pupil dynamics^30^—and extend them to a functional network level.

**Figure 4.**
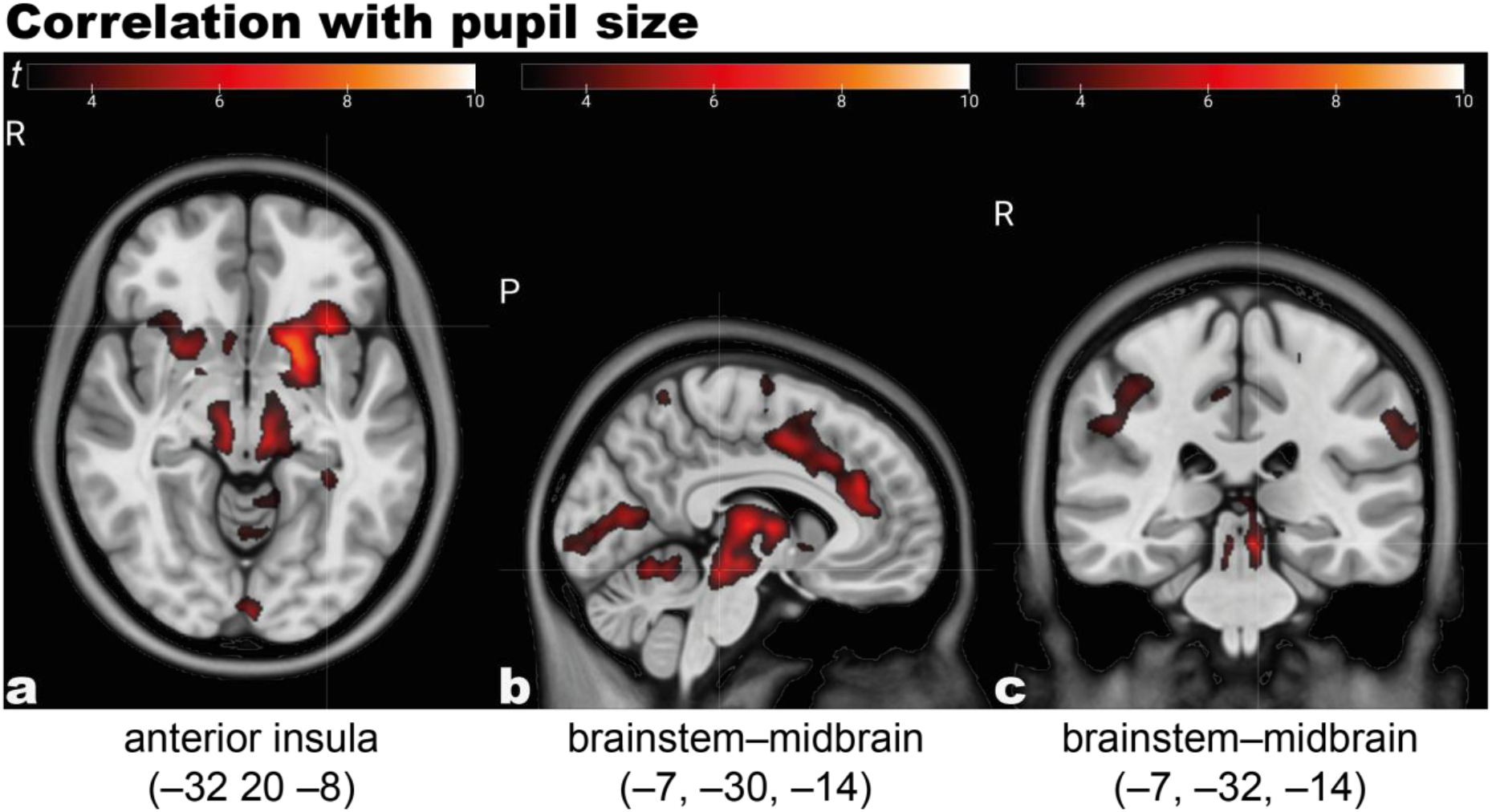
Moment-to-moment changes in pupil size correlate with action-mode network (a) and neuromodulatory brainstem–midbrain activation (b–c). For visualization, group contrast maps are thresholded at *t =* 3, main text results report family-wise error corrected statistics. Contrast maps are available at: osf.io/4t5hj. Numbers below the labeled regions indicate MNI XYZ-coordinates.

Importantly, these pupil–fMRI analyses also revealed a bilateral brainstem–midbrain cluster (*t =* 7.27, *p*_FWE-corr_ < 0.001 MNI: –4, –30, –14), overlapping with key noradrenergic, serotonergic, and dopaminergic nuclei, in accordance with recent work in rodents^55,65,66^ and primates^67–69^. See Figure S3 for analyses probing the quality of brainstem alignment and Figures S4–5 as well as Table S1 for locus coeruleus-specific pupil–BOLD associations.

In sum, we found that neuromodulatory and action-mode network regions activated when pupils dilated, suggesting neuromodulatory drive of network transitions. For inverse associations of moment-to-moment pupil and locus coeruleus signal changes with default-mode network activity, see Table S2–3.

### Noradrenergic transporter distribution overlaps with pupil-linked BOLD activation patterns

If indeed the observed activation pattern is supported by noradrenergic release in the cortex, one would expect overlapping spatial distributions with the noradrenergic system. To test this, we leveraged public PET-derived maps of the cortical and subcortical distribution of the noradrenergic transporter^56,70–74^. We then assessed the voxel-wise association of noradrenergic transporter expression and pupil-linked BOLD activation. We detected a robust correlation (Hesse et al. map: *r* = 0.189; *p*_permutation-corr_ < 0.001), that we replicated using an alternative noradrenergic transporter map (Ding et al. map: *r* = 0.161; *p*_permutation-corr_ < 0.001; Figure 5). This suggests that those parts of the brain that activated when pupils dilated overlapped with those housing noradrenergic receptors.

**Figure 5.**
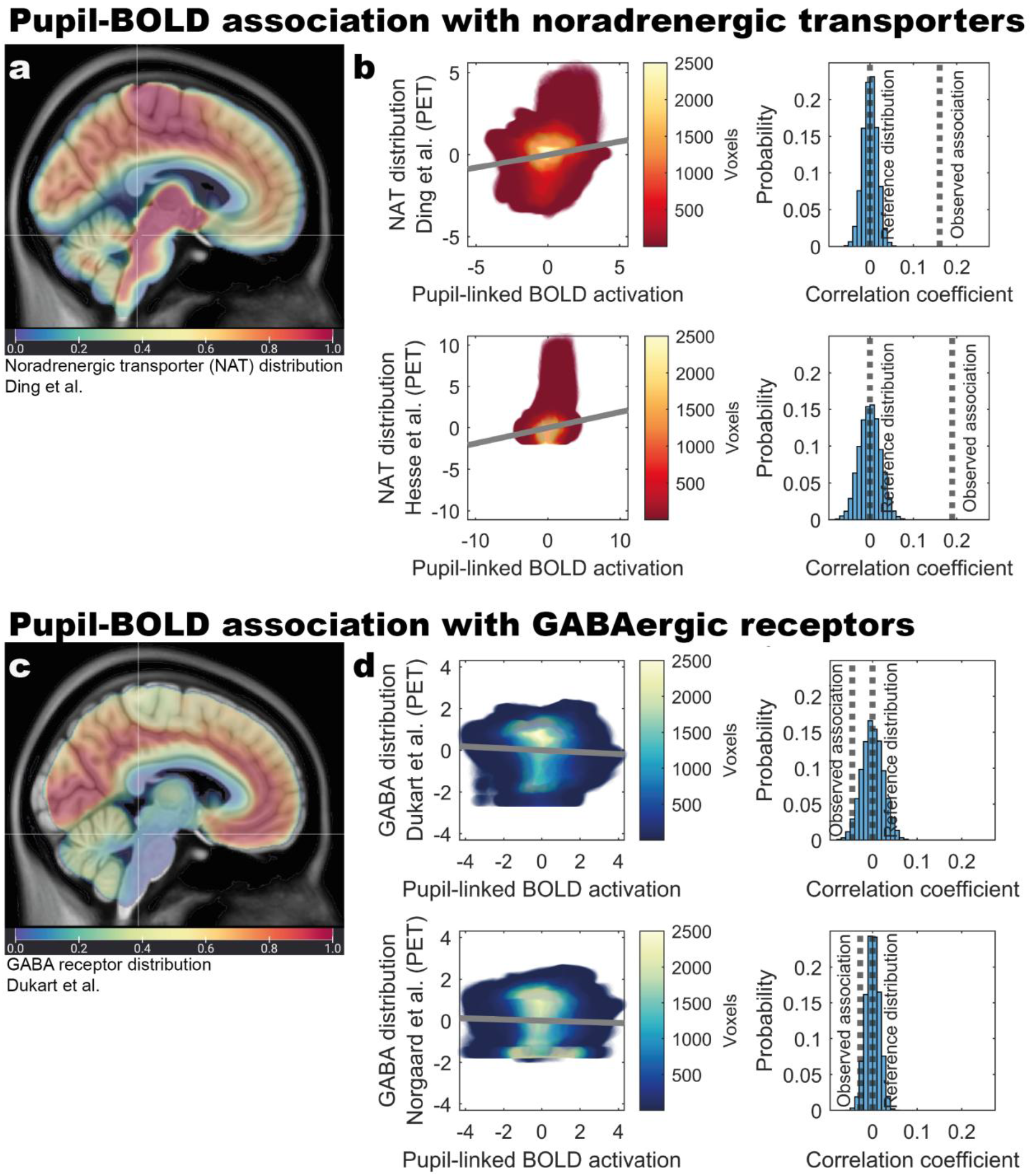
Voxel-wise association of pupil-linked BOLD activation patterns with PET-derived maps of noradrenergic transporter (a–b) and GABAergic receptor (c–d) distribution. (**a**) Relative distribution of noradrenergic transporters based on^74^. (**b**) Brain regions activating when pupils dilate overlap with noradrenergic transporter distribution based on two alternative PET maps. (c) Relative distribution of GABA_A_ receptors based on^75^. (**d**) Brain regions activating when pupils dilate show no systematic overlap with GABAergic receptor distribution based on two alternative PET maps. Observed correlation coefficients are tested against a reference distribution obtained using phase-randomization permutation.

To further contextualize these results, we computed two negative control analyses. Specifically, we linked two brain-wide GABA_A_ receptor distribution maps^75,76^ to our pupil-linked BOLD maps, which we assumed to show no or even a negative association^77^. In line with this, we found that GABA_A_ receptor expression showed a non-significant negative relation with the pupil–fMRI pattern (Nørgaard et al map: *r* = –0.028; *p*_permutation-corr_ = 0.063; Dukart et al. map: *r* = –0.045; *p*_permutation-corr_ = 0.056; Figure 5).

Finally, given that large parts of the brainstem–midbrain, overlapping with noradrenergic, but also serotonergic and dopaminergic nuclei, activated when pupils dilated, we ran two additional analyses to determine whether similar effects extended to other neuromodulatory systems. Individually, both serotonergic (Fazio et al. map: *r* = 0.246; *p*_permutation-corr_ < 0.001;^56^) and dopaminergic (Sasaki et al. map: *r* = 0.241; *p*_permutation-corr_ < 0.001;^78^) transporter expression showed a comparable spatial correspondence with the pupil–fMRI activation pattern. While the cortical innervation of dopaminergic, serotonergic and noradrenergic nuclei is partially overlapping, there are also considerable differences between these neuromodulatory systems^3^. To test if the expression pattern of noradrenergic transporters explains unique variance in the observed pupil-linked BOLD maps over and above the other neuromodulatory systems, we estimated multiple regression analyses. Specifically, we compared a base model (including serotonergic and dopaminergic transporter expression as predictors) to a full model additionally including a predictor for the noradrenergic transporter (Table 1). Importantly, we found the full model substantially outperformed the base model, suggesting a unique association of noradrenergic transporter expression and pupil-linked BOLD activation (Δ*χ*²(*df* = 1) = 27373; *p*_permutation-corr_ < 0.001; for the explained variance of both models, see Table S4). In addition to voxel-wise spatial correlations (Figure 5, Table 1), we quantified overlap using Dice coefficients (Figure 6, Table 2), yielding comparable results.

**Figure 6.**
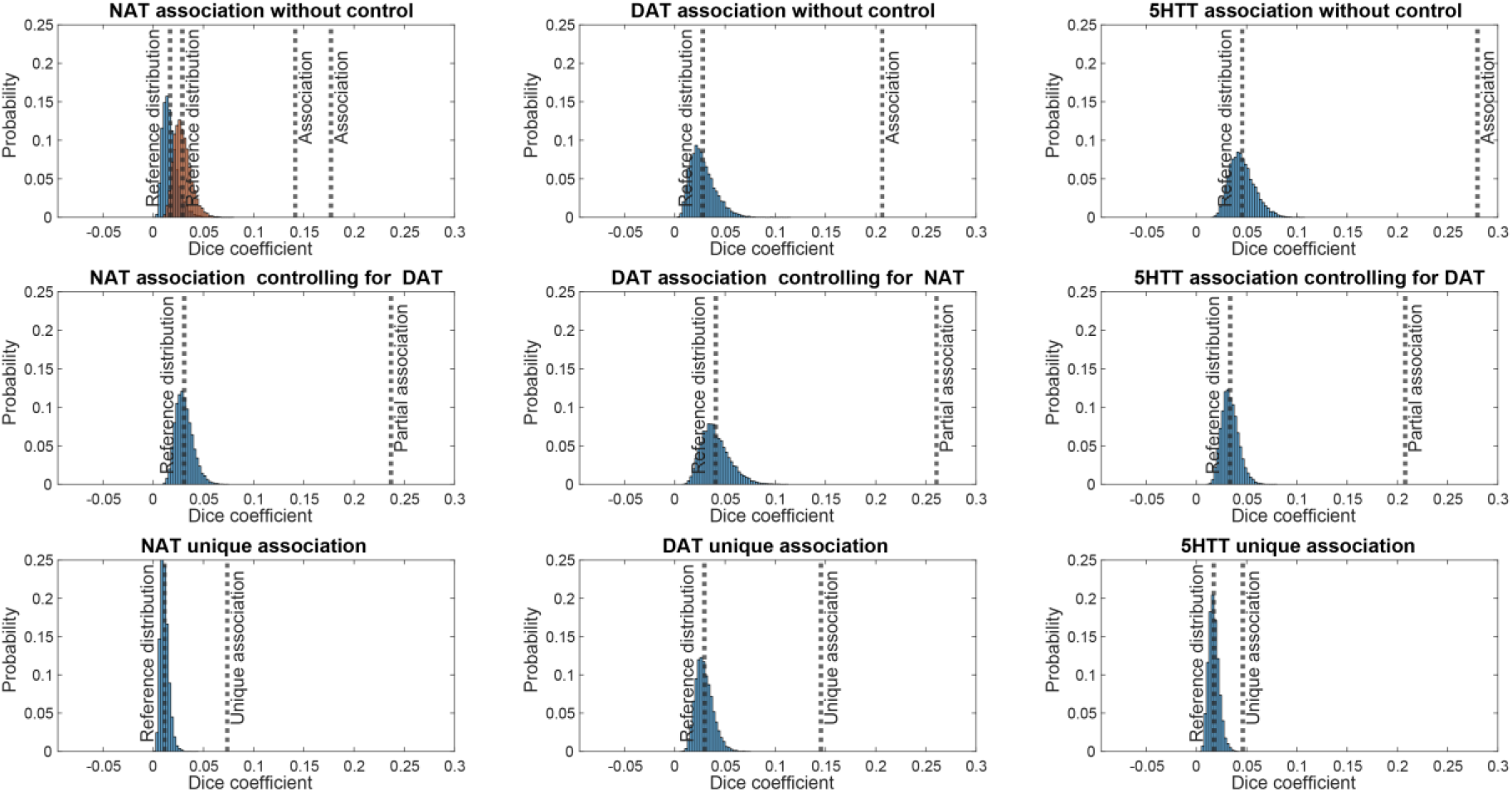
Spatial correspondence between pupil-linked BOLD activity and neuromodulatory transporter distributions. Dice coefficients quantify the overlap between the standardized and binarized z (≥ 1.96) group-level pupil–BOLD activation map and PET-derived transporter maps (noradrenergic, NAT; dopaminergic, DAT; serotonergic, 5-HTT). Observed associations are tested against a reference distribution obtained using phase-randomization permutation. Partial Dice coefficients estimate neuromodulatory system-specific overlap after excluding voxels shared with other transporter maps. The Ding et al. and Hesse et al. NAT maps^79,80^ were averaged for estimations of partial associations (middle and bottom row), while the first row shows their individual associations (dotted lines) and corresponding reference distributions (blue, orange).

**Table 1:** Multiple regression predicting pupil-linked BOLD activation based on neuromodulatory transporter maps.

| Predictor | Estimate | Standard error | t-value | p |
| --- | --- | --- | --- | --- |
| NAT | 0.1780297 | 0.001071632 | 166.12949 | <0.001 |
| 5HTT | 0.071771108 | 0.001026131 | 69.943398 | <0.001 |
| DAT | 0.21053788 | 0.000838879 | 250.97537 | <0.001 |

Note: NAT, 5HTT, DAT refer to PET-derived noradrenergic, serotonergic and dopaminergic transporter expression maps. To obtain a single noradrenergic predictor, the Hesse et al. and Ding et al. maps were averaged.

**Table 2:** Raw and corrected Dice coefficients expressing spatial overlap between pupil–BOLD and neuromodulatory transporter maps.

| PET transporter map | Raw (partial)<br>Dice coefficient | Reference distribution<br>mean and standard deviation |  | Corrected (partial)<br>Dice coefficient |
| --- | --- | --- | --- | --- |
| NAT (Ding et al.) | 0.141 | 0.017 | 0.008 | **8.37** |
| NAT (Hesse et al.) | 0.177 | 0.029 | 0.008 | 6.108 |
| 5HTT (Fazio et al.) | 0.28 | 0.046 | 0.012 | 6.138 |
| DAT (Sasaki et al.) | 0.206 | 0.028 | 0.012 | 7.407 |
| NAT controlling for DAT | 0.237 | 0.031 | 0.009 | **7.652** |
| DAT controlling for NAT | 0.261 | 0.041 | 0.014 | 6.403 |
| 5HTT controlling for DAT | 0.208 | 0.034 | 0.008 | 6.191 |
| NAT unique | 0.074 | 0.011 | 0.004 | **6.535** |
| DAT unique | 0.145 | 0.029 | 0.009 | 4.944 |
| 5HTT unique | 0.046 | 0.017 | 0.005 | 2.661 |

*Note:* Dice coefficients quantify the overlap between the standardized and binarized $z$ ( $\geq 1.96$ ) group-level pupil–BOLD activation map and PET-derived transporter maps (NAT, DAT, 5-HTT). Partial Dice coefficients estimate neuromodulatory system-specific overlap after excluding voxels shared with other transporter maps. Observed associations are tested against a reference distribution obtained using phase-randomization permutation. Corrected Dice overlap coefficients are estimated by normalizing observed Dice coefficients relative to the mean of the corresponding null distribution (i.e., the strength of observations expected by chance). Values highlighted in bold indicate the winning transporter map for each comparison.

Taken together, these analyses suggest that key ascending arousal-promoting systems, including noradrenergic neuromodulation, contribute to the observed pupil-linked subcortical and action-mode network activation.

### Locus coeruleus is functionally connected to action-mode network hubs

So far, our analyses suggest that pupil dilation, likely reflecting neuromodulatory drive, is associated with brainstem and action-mode-network activations. Moreover, the activated regions closely overlap with noradrenergic receptor distributions as derived from PET atlases. These observations prompted us to test if locus coeruleus and action-mode network activation were directly connected.

Using dedicated neuromodulation-sensitive MRI^49^, we estimated person-level proxies for locus coeruleus integrity as well as a group image that visualizes the spatial location of the locus coeruleus adjacent to the fourth ventricle (Figure 7 and osf.io/4t5hj). Notably, the locus coeruleus-related hyperintensity was adequately captured by a previously generated high-confidence mask of this structure^81^ (available at: https://osf.io/sf2ky/).

**Figure 7.**
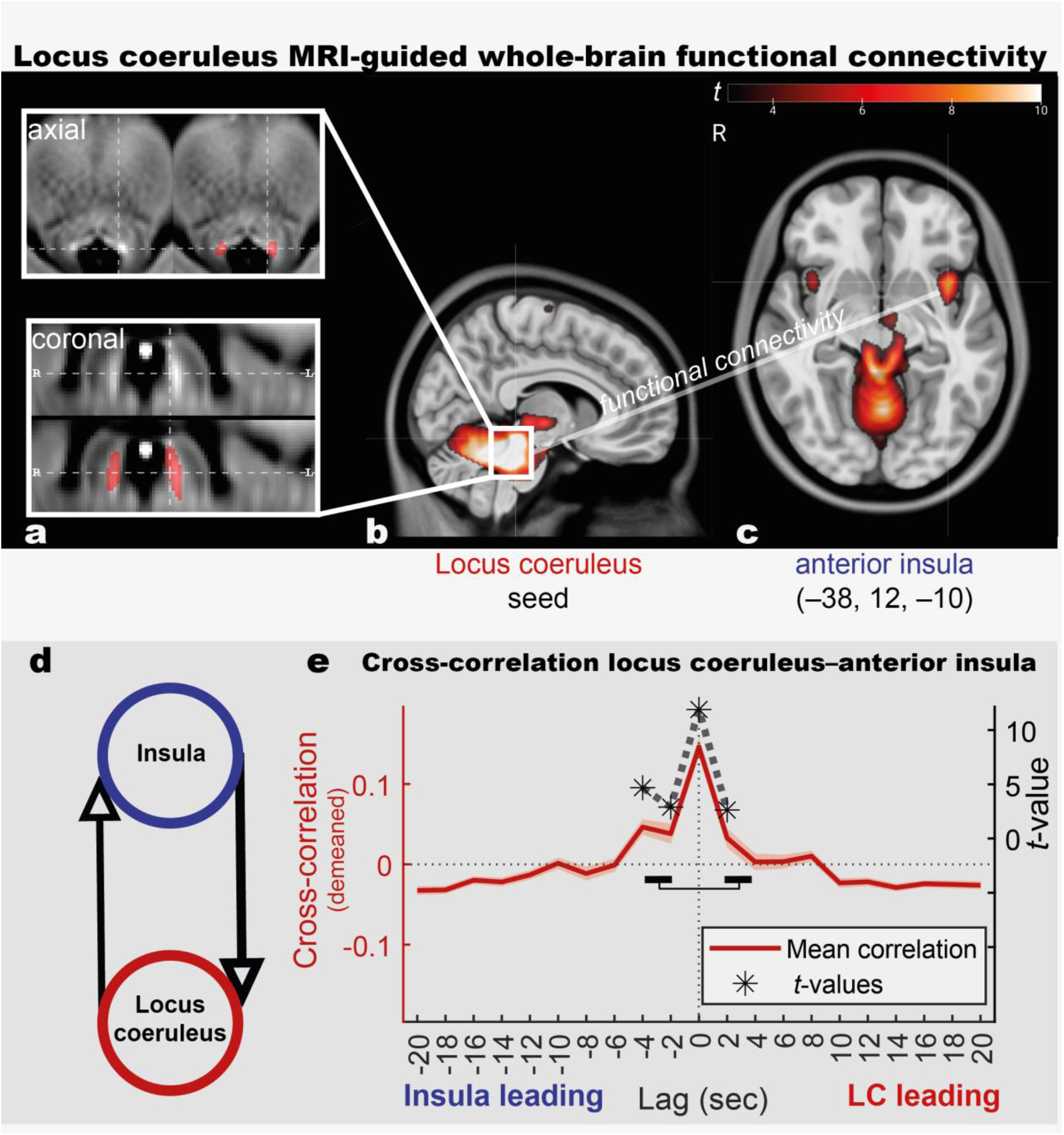
Locus coeruleus MRI (a) guides whole-brain functional connectivity analyses (b–c), revealing action-mode network activity is coupled to and precedes brainstem activation (d–e). (**a**) Neuromodulation-sensitive MRI reveals locus coeruleus-related hyperintensity bordering the fourth ventricle that is accurately captured by a high-confidence volume of interest (red;^81^). (**b**) Using this region as seed for whole-brain task-based functional connectivity shows locus coeruleus–anterior insula co-fluctuations. (**d–e**) Cross-correlations again show a locus coeruleus–anterior insula association and suggest this is initiated by the anterior insula, showing a temporal precedence of up to 4 s (contrast of lags [(–4)–(–2)] vs [(+2)–(+4)]). In panel (**e**), the dark red line represents the mean cross-correlation coefficients, averaged across participants, while the lighter red shaded line represents the standard error of the mean. The dashed grey line indicates the corresponding *t*-values, resulting from a group-level cluster-based permutation test evaluating the significance of the cross-correlation for each sample. For additional Dynamic Causal Modeling-based locus coeruleus–anterior insula effective connectivity analyses^85,86^, see Figures S6–7.

After this validation step, we used the derived coordinates overlapping with the masked area as seed region for trial-level whole-brain functional connectivity analyses, probing which regions’ activity co-fluctuated with the locus coeruleus during the conditioned oddball task^82,83^. A bilateral anterior insula (*t =* 8.93, *p*_FWE-corr_ < 0.001 MNI: –38 12 –10; Figure 7) and thalamus (*t =* 6.79, *p*_FWE-corr_ < 0.001 MNI: –6.5 –22.5 –2.5) cluster survived correction for multiple comparisons, in addition to a broad cerebellar and pontine activation. The un/thresholded whole-brain contrast maps are available at osf.io/4t5hj. These results point to a co-fluctuation of activity in the locus coeruleus and key action-mode network hubs, in line with known direct anatomical projections^9^, as well as our pupil–fMRI and noradrenergic transporter findings. We repeated our functional connectivity analyses^83^ using minimally smoothed locus coeruleus BOLD data as seed input (2 mm FWHM). For the primarily cortical target voxels, we maintained the default smoothing (8 mm FWHM). Replicating our earlier results, we again found a left anterior insula cluster (MNI: –38 –14 –10) co-fluctuating with locus coeruleus activity (*t* = 6.4, *p*_FWE-corr_ < 0.001). For inverse associations of locus coeruleus and default mode network activity, see Table S3.

However, the functional connectivity analyses do not speak to the directionality of this association (brainstem→action-mode network vs. action-mode network→brainstem). Thus, we ran a separate set of analyses, using the TR-level BOLD timeseries of the locus coeruleus and anterior insula (i.e., on a sample level, averaged over voxels). We submitted these to a cross-correlation analysis, which measures the temporal relationship between two signals by quantifying how one variable aligns with shifts in another over time^84^, allowing us to assess the temporal precedence of brainstem vs. action-mode network activation. First, we observed a strong coinciding activation (mean cross-correlation coefficient at lag zero: 0.363; *t =* 11.892), in line with our functional connectivity findings, that stood out above the general association of the two signals (mean cross-correlation coefficient across all lags: 0.217). Note that to better judge the temporal relation of activations, we removed this overall background association by demeaning before group-level cluster-statistics (Figure 7).

Adding to our connectivity analyses, we found that anterior insula activations preceded and predicted subsequent locus coeruleus activations by up to four seconds (mean cross-correlation coefficient at lag –4 s: 0.264; *t =* 4.69; at lag –2 s: 0.255; *t =* 2.896; cluster permutation test: *t*_sum_ = 22.103; *p*_cluster-corr_ ≤ 0.002; cluster including lags: (–4)–(+2) s). We also observed that preceding locus coeruleus activity was followed by action mode network activation two seconds later (mean cross-correlation coefficient at lag +2 s: 0.25; *t =* 2.626), suggesting a potentially slightly later recurrent connection (brainstem↔action-mode network; contrast of cross-correlation coefficient at lags [–4 to –2; anterior insula leading] vs [+2 to +4; locus coeruleus leading]: *t =* 2.421; *p* = 0.018). While these analyses do not allow for causal interpretations, they indicate that anterior insula activity may recruit noradrenergic neuromodulation to facilitate subsequent action-mode network activation^5,6^.

For additional Dynamic Causal Modeling-based locus coeruleus–anterior insula effective connectivity analyses^85,86^ that support this interpretation, see Figures S6–7 and Table S5.

Taken together, we observed that brainstem activation was functionally connected with action-mode network activity and that this crosstalk may be initiated by the anterior insula, potentially helping participants to efficiently respond to unexpected stimuli^2^.

### Locus coeruleus and action-mode network support implicit learning of task structure

To investigate the implications of locus coeruleus–action-mode network interactions during the conditioned oddball task, we analyzed behavioral data. As participants showed high overall accuracy (group mean >0.95), our analyses focused on response time dynamics. Participants entered the task without prior knowledge of its statistical structure, yet mean response times systematically varied with trial likelihood across the fMRI session (Figure 8a; *F* = 173.83; *p* < 0.001; mean response times per condition: stay-standard: –0.437±0.021, switch-to standard: 0.173±0.031, stay-oddball: 0.442±0.043, switch-to oddball: 0.602±0.040; note that all response times are expressed in *z*-values after log-transformation and within-person standardization). This pattern suggests that participants responded quicker to frequent trial types at the cost of slower responses to rare events. Supporting this, a strong negative association of the response times for stay-standard and switch-to-oddball conditions indicates a performance trade-off between these categories (Figure 8b; *r* =–0.795; *p* < 0.001). We thus assumed that participants implicitly learned the statistical structure of the novel task environment and adapted their behavior, which implies that this response time pattern should emerge over the course of the experiment. Combining behavioral data from both repetitions of the conditioned oddball task, we observed that early in the experiment, responses were generally slow and did not differentiate between frequent and rare trials (Figure 8c). However, as the experiment progressed, response times to common stimuli became faster, while rare oddballs elicited increasingly slower responses. Accordingly, we found that the difference in response times between switch-to-oddball and stay-standard categories increased with time, suggesting implicit learning (Figure 8d; correlation of response-time difference and time bin on a group level: *r* = 0.509; *p* = 0.003). This learning pattern detected on a group level was also evident within participants (dependent samples *t*-test of within-participant time-bin beta parameters against zero: *t =* 3.194, *p* = 0.002; mean beta parameter: 0.0097±0.00035; Figure S8). Collectively, these results indicate that participants, initially unfamiliar with the task, gradually internalized its structure and adjusted their behavior accordingly. For additional analyses of response time dynamics over the course of the conditioned oddball task, including trial-level linear mixed effects models, see Table S6 and Figure S9.

**Figure 8.**
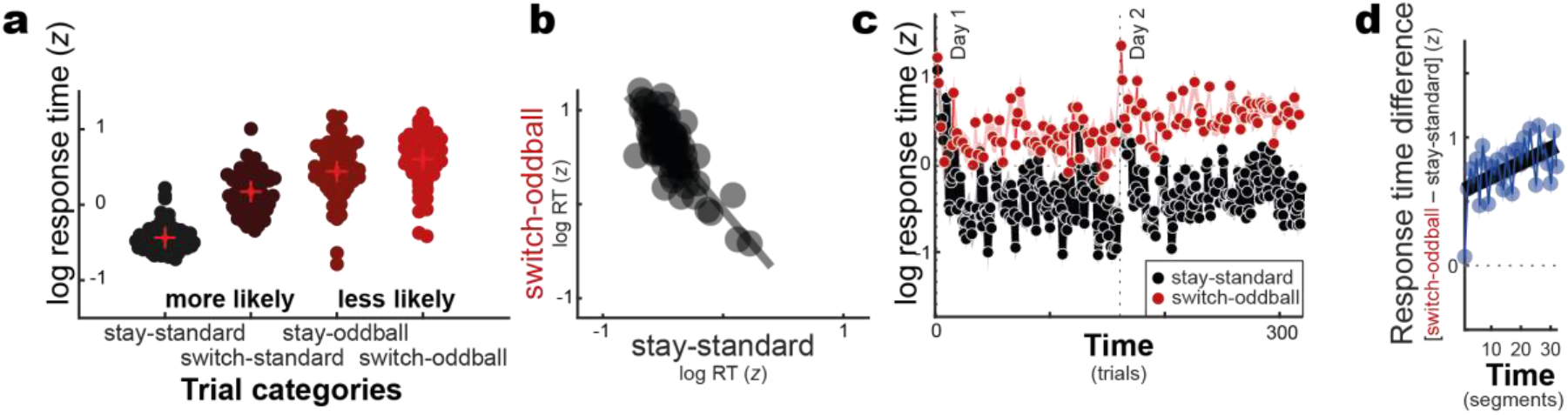
Participants focus their behavior on more likely trial categories (a) at the expense of less likely categories (b). This bias emerges across the course of the experiments (c) indicating implicit learning of the task structure (d). Day 1 indicates the pupillometry-EEG assessment, while day 2 indicates the pupillometry-fMRI oddball assessment. (**a–b**) show day 2 data. (**c–d**) include data of both days.

Finally, we combined imaging and behavioral data to probe if individual differences in markers of the noradrenergic system and its cortical target regions explained implicit learning performance. Specifically, we combined dedicated structural locus coeruleus imaging (Figure 7a and Figure 9a left), task-related pupil dilation (cf. Figure 2b) and locus coeruleus-related anterior insula activation (cf. Figure 3a) in a multivariate partial least squares correlation with condition-wise response times (stay-standard; switch-to standard; stay-oddball; switch-to oddball; Figure 8 and Figure 9a right). We identified one reliable latent component (*r* = 0.488; *p*_permutation-corrected_ = 0.001). Participants with stronger task-related anterior insula activation (loading onto latent variable: 0.752; bootstrap ratio: 3.402 [interpreted analogous to *z*-values]), higher locus coeruleus integrity (loading: 0.491; bootstrap ratio: 2.306) and task-related pupil dilation (loading: 0.44; bootstrap ratio: 1.873; trend level association) showed more pronounced differentiation in response times between frequent and rare stimuli (cf. Figure 9b). In particular, these participants tended to show quicker responses for frequent trial categories (loadings; stay-standard: –0.468; switch-to-standard: –0.396) at the expense of slower responses for infrequent trial categories (loadings; stay-oddball: 0.364, switch-to-oddball: 0.701)—recapitulating the implicit-learning pattern we have identified in behavioral analyses (Figure 8c). Brain–behavior analyses were evaluated in a multivariate statistical framework^87^, given its higher sensitivity, followed up by post-hoc tests split by neural indicator (insula BOLD: *r* = 0.356, *p* = 0.003; locus coeruleus integrity *r* = 0.224, *p* = 0.067; pupil dilation: *r* = 0.207, *p* = 0.090; also see Table S8). For analyses split by trial category, see Figure S10 and Table S7.

**Figure 9.**
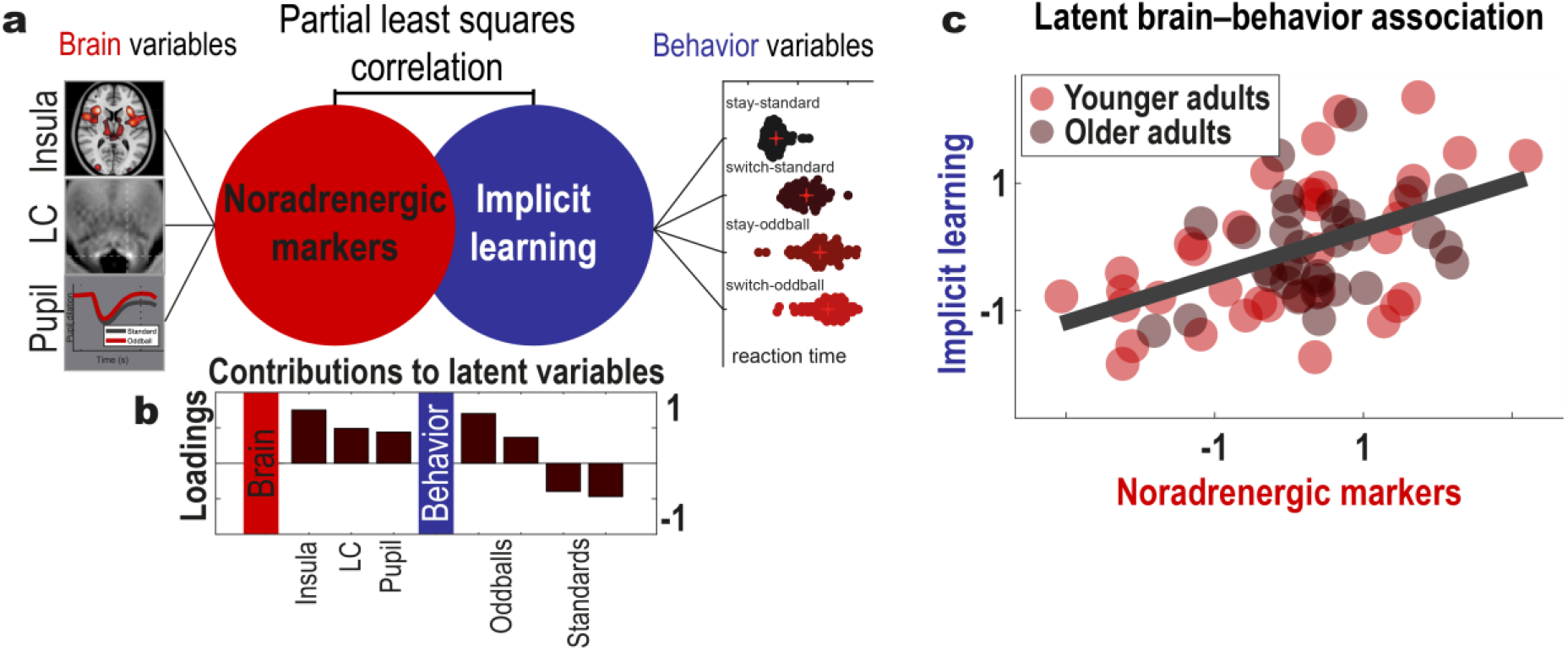
Individual differences in noradrenergic markers (insula BOLD activity, locus coeruleus integrity, pupil dilation) are associated with implicit learning. (**a**) Multivariate partial least squares correlation links noradrenergic markers to response times for frequent and infrequent trial categories (standards, oddballs, respectively), which reflect that participants implicitly learned the statistical regularities of the task. (**b**) Noradrenergic markers positively contribute to the latent brain variable (e.g., the more activation, the higher scores). On a behavioral level, quicker response times for frequent trial categories at the expense of slower response times for rare categories are linked to higher latent behavioral scores. (**c**) Participants with higher values in markers of the noradrenergic system demonstrate more implicit learning, a pattern that is comparable across age groups.

Post-hoc age group comparisons revealed that younger and older adults showed comparable latent behavioral (*t =* –1.36; *p* = 0.179) and neural scores (*t =* –0.209; *p* = 0.835) and that the latent association did not differ across groups (*z =* 1.111; *p* = 0.133; cf. Figure 9c and Table S9 in the Supplementary Information).

As a sensitivity analysis, we replicated partial least squares correlation findings using a simpler analytical strategy (*r* = 0.466; *p* < 0.001; by averaging over standardized neural indicators (i.e., insula BOLD activity, locus coeruleus integrity, and pupil dilation, see PLSC analyses) and correlating this composite score with the difference in response times to oddball and standard stimuli).

In summary, participants adapted their responses over time to reflect the statistical regularities of the experimental task, demonstrating implicit learning. This implicit learning pattern was more strongly expressed in those participants with higher insula activation, higher structural locus coeruleus integrity, and a stronger pupil response to salient stimuli. Together those findings collectively support a role of the locus coeruleus-noradrenergic system in guiding attention and learning to adapt behavior in novel environments^3,6,41^.

### Conditioned oddball stimuli elicit pupil-linked P300 and cortical excitability modulations

Prior to the pupillometry–fMRI session (result Figures 3–9), the same participants completed the conditioned oddball task while we simultaneously recorded pupil and EEG activity (cf. Figure 1–2). This allowed us to examine complementary electrophysiological signatures of noradrenergic activation^4,57,58,61^ and to what extent these covary with pupil-indexed neuromodulation.

Specifically, previous work suggests a positive parietal event-related potential around 300 ms after stimulus onset—the P300—is influenced by phasic locus coeruleus activity and supports the cortical encoding of salient information^57,58^. In line with this, relative to standard stimuli, oddballs elicited a significantly more robust P300 potential (*p*_cluster-corr_ ≤ 0.002; Figure 10a), which positively scaled with changes in pupil-linked neuromodulation on a trial level (*p*_cluster-corr_ ≤ 0.036; Figure 10b; for the corresponding pupil results, see Figure 2b).

**Figure 10.**
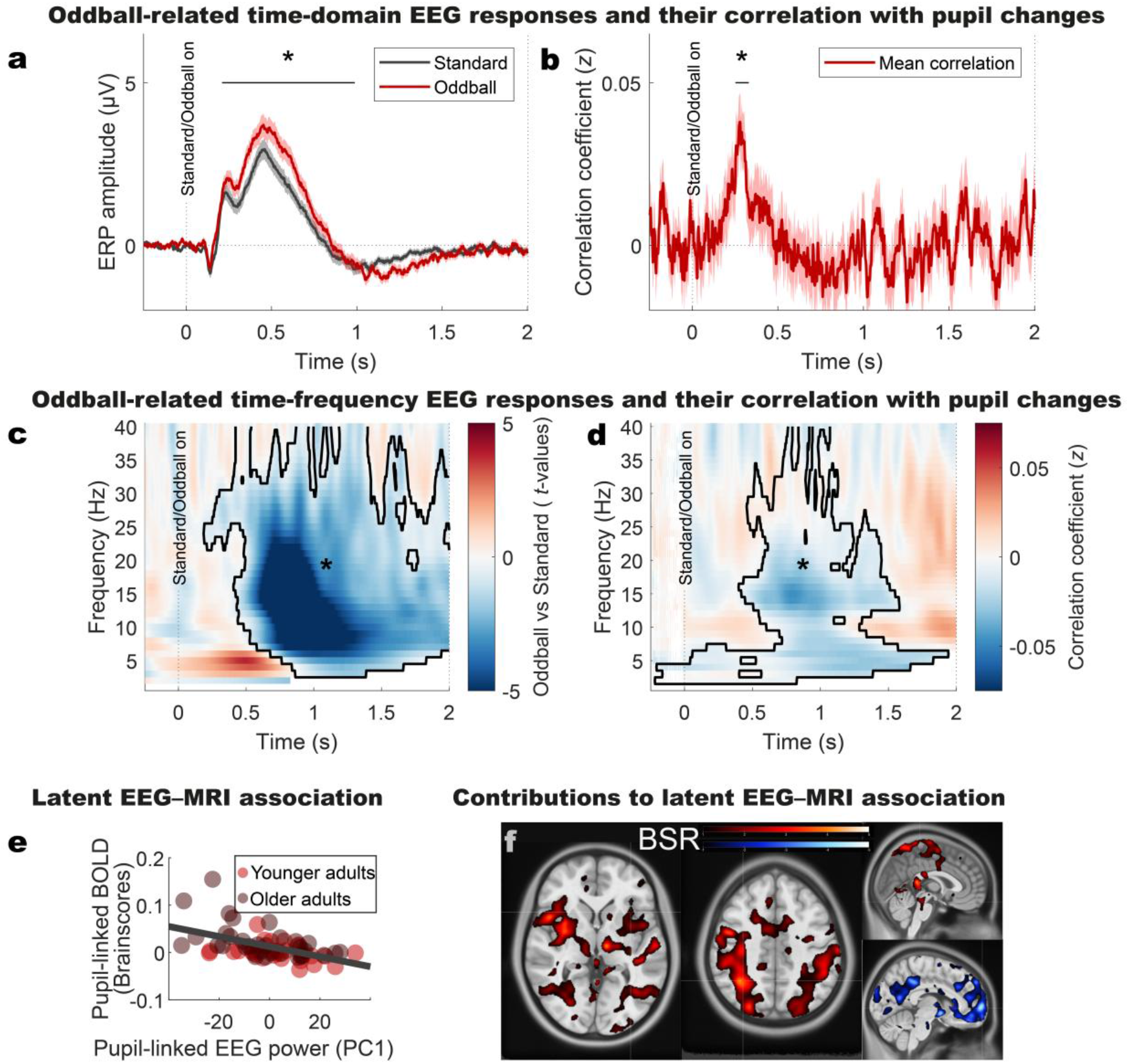
Salient oddball stimuli elicit electrophysiological time-domain (a–b; P300 event related potential) and time-frequency domain (c–d; low frequency power suppression) correlates of neuromodulatory activation that are linked to pupil-linked BOLD modulations (e–f). **(a)** Oddball (red) relative to perceptually matched Standard stimuli (black) demonstrate a more positive event-related potential overlapping in time with the P300, an established electrophysiological marker of noradrenergic activity^98,99^. **(b)** In line with its association with neuromodulation, trials with higher amplitudes within the P300 time window show greater pupil dilation, leading to a positive correlation at the trial level (red). Correlations were estimated across all trial categories. **(c)** Oddball relative to standard stimuli also demonstrate a pronounced suppression of low-frequency EEG power, an electrophysiological marker for cortical excitability related to noradrenergic activity^100,101^. **(d)** Trials showing more suppression of low-frequency EEG power also show greater pupil dilation, suggesting an association with neuromodulation. Correlations were estimated across all trial categories. **(a–d)** For visualization, EEG signals are averaged across electrodes identified by group-level cluster-permutation tests. Pupillometry–EEG correlations reflect within-participant trial-by-trial associations. Shaded areas denote ± standard error of the mean. Asterisks and contours indicate significant clusters. Only the primary clusters are reported **(e–f)** Association between pupil-linked EEG power (first principal component scores; PC1) and pupil-linked BOLD brain scores across participants were derived using partial least-squares correlation (PLSC). **(e)** The between-participant latent-level correlation of pupil-linked EEG power and pupil-linked BOLD activity is shown with marker color indicating the age group. Participants showing more negative pupil-linked EEG power (cf. Figure 10c) also show more positive pupil-linked BOLD activity (cf. Figure 4). **(f)** Voxel-wise contributions to the latent EEG–fMRI association are quantified by bootstrap ratios (BSR). Regions with positive and negative contributions are shown in warm and cool colors, respectively. Overlap with canonical functional networks is summarized in Tables S10–11.

Cortical excitability fluctuates during wakefulness^88^, with noradrenaline release promoting states of higher excitability^4,59–61,89,90^ that facilitate the processing of incoming information^91,92^. Accordingly, oddball stimuli led to pronounced suppression of low-frequency EEG power, most pronounced in the alpha–beta bands (∼8–20 Hz), an established electrophysiological marker for cortical excitability^4,39,93–95^ (*p*_cluster-corr_ ≤ 0.002; Figure 10c). Trials involving stronger decreases in low-frequency power (i.e., an increase in cortical excitability) were accompanied by larger pupil dilation, pointing to neuromodulatory contributions^4^ (*p*_cluster-corr_ ≤ 0.006; Figure 10d). If pupil-linked electrophysiological cortical excitability markers capture functionally relevant downstream outcomes of noradrenergic engagement, these should be associated with implicit learning during the conditioned oddball task, similar to the MRI markers (cf. Figure 9). Indeed, we observed that participants with stronger pupil-linked excitability modulations (i.e., more low-frequency EEG power decreases) demonstrated better adaption to the statistical structure of the task, consistent with enhanced implicit learning (*r* = –0.269; *p* = 0.021; see Figure S11 for how the dimensionality of the EEG data was reduced for these analyses).

Finally, we used pupil tracking as a physiological bridge between the two assessment days (cf. Figure 1) and the EEG and fMRI modalities to assess common neuromodulation-linked mechanisms. Participants who showed stronger pupil-linked excitability modulations (i.e., low-frequency EEG power suppression) also showed stronger pupil-linked BOLD modulations (Partial Least Squares-derived latent-space association: *r* = −0.522, *p*_permutation-corrected_ = 0.031; Figure 10e).

Incorporating spatial information revealed that positive contributors to this association were primarily located in the action-mode network, dorsal attention network, as well as somato–motor and brainstem–midbrain regions (Table S10; Figure 10f), whereas the default-mode network showed the opposite relationship (i.e, deactivation; Table S11; Figure 10f). Together, these findings suggest that pupil-indexed neuromodulatory responses are accompanied by coordinated changes in cortical excitability and large-scale functional network transitions across measurement modalities^4–6,96,97^.

## 3. Discussion

This study investigated the role of the locus coeruleus and its cortical targets in how younger and older participants adapt their behavior in dynamic environments with novel observations. We took advantage of repeated assessments of an experimental task shown to modulate noradrenergic activity in primates^44^ and a dedicated multimodal imaging protocol to identify neuromodulatory contributions to implicit learning.

We found that salient expectation-violating stimuli dilated participants’ pupils, indicating heightened neuromodulatory activity, and triggered a shift from default-mode to action-mode network activation that was associated with increased cortical excitability. Combining pupil and BOLD data, we observed that neuromodulatory brainstem–midbrain as well as action-mode network regions scaled their activity with pupil size. This suggests an interplay between the noradrenergic system and the action-mode network, which we confirmed using PET-derived transporter maps and locus coeruleus-centered functional connectivity analyses. Behavioral data showed that participants implicitly learned the statistical structure of the novel task environment, resulting in quicker responses for more prevalent trial categories at the expense of slower responses to rare oddballs. Linking brain and behavior, we found that this implicit learning pattern was more pronounced in participants with higher levels of markers associated with the noradrenergic system and its cortical targets. Taken together, this study suggests noradrenergic neuromodulation may upregulate the action-mode network to facilitate (implicit) learning for behavioral adaptation.

### Fear conditioned stimuli evoke robust neuromodulatory responses

To increase recruitment of the locus coeruleus, a subset of our experimental stimuli was pre-conditioned before the main task by repeatedly pairing them with an aversive outcome^40,44–46^. Conditioned stimuli (CS+) elicited greater pupil dilation than perceptually matched control stimuli (CS–), suggesting increased neuromodulatory drive. Moreover, these effects remained stable across age groups and assessment days, highlighting the robustness of the conditioned response. Our findings are in line with earlier work in rodents demonstrating increased locus coeruleus spiking^102^ and noradrenergic axonal activation^45^ during conditioning, assumed to support fear learning.

### Noradrenergic neuromodulatory activation is coupled with the action-mode network

Building on this, during the conditioned oddball task, salient stimuli dilated participants’ pupils, enhanced the P300 event-related potential, and activated the action-mode network, including the anterior insula, middle cingulate cortex, thalamus, and cerebellum. This aligns with prior findings implicating the brain’s action-mode of function with states of externally focused attention, heightened arousal, and the processing of action-relevant bottom up signals^22,25^. Importantly, these activations were accompanied by default-mode network deactivation, suggesting a functional transition from introspective to externally oriented processing^6^, which on an electrophysiological level was coupled to increased cortical excitability^4^. This is in line with the proposed antagonism of the default-mode and action-mode networks^22^ and the more general noradrenergic role in facilitating dynamic reorganizations of target neural networks (also termed network reset^5^).

Combined analyses of pupil and BOLD time series revealed that moment-to-moment fluctuations in pupil size systematically co-varied with activation in the action-mode network (anterior insula, middle cingulate cortex, thalamus). This supports previous intracranial work in humans linking anterior insula activations to variations in pupil size^30^ and extends them to a network level. In addition, a brainstem–midbrain cluster encompassing the locus coeruleus and other neuromodulatory nuclei activated when pupils dilated, in line with recent animal research^55,65–67^, suggesting that pupil dilation reflects broad neuromodulatory engagement across cortical and subcortical regions^68,69^. However, while useful for constraining the hypothesis space, our current correlation-based analysis approach is ambiguous regarding the directness of this association^103^ (but see Figure S5).

An analysis of the spatial overlap between publicly available PET-derived noradrenergic transporter maps and pupil-linked BOLD activation further underscores a role for noradrenergic neuromodulation in regulating these activation patterns^56,70–74^. We found a robust positive correlation, replicating across independent transporter maps, suggesting that brain regions with increased activity during pupil dilation overlap with areas rich in noradrenergic transporters. Negative control analyses with GABAergic receptor maps^75,76^ showed no such association, further strengthening the specificity of the association between pupil size, locus coeruleus and action-mode network activation. In line with the broad brainstem–midbrain activation, also serotonergic^104^ and dopaminergic^78^ transporter maps were positively coupled with the pupil-linked BOLD pattern^68,69^. Crucially, however, noradrenergic transporter expression explained unique variance in pupil-linked BOLD activation over and above the other neuromodulatory systems.

To more precisely map locus coeruleus activation to its cortical targets, we employed functional connectivity analyses guided by dedicated neuromodulation-sensitive MRI^81^. These analyses revealed that locus coeruleus activity co-fluctuated with action-mode network hubs, particularly the anterior insula and thalamus, consistent with established direct projections in primates^9^. Cross-correlation analyses further elucidated the temporal dynamics of this relationship, suggesting that the anterior insula may recruit locus coeruleus activation up to four seconds later. This finding together with evidence of a recurrent connection support the notion that the insula-initiated recruitment of noradrenergic signaling may facilitate a cortical shift toward an action-oriented state, optimizing adaptive responses to environmental changes^5,6,22^. In this circuit, the anterior insula likely provides crucial contextual signals without which the locus coeruleus would be blind to salient events that require gain-modulation^2^. Our findings are in broad agreement with a recent rodent study in which direct stimulation of the cingulate cortex, another action-mode network hub region, elicited pupil dilation through locus coeruleus activation, suggesting top-down recruitment of brainstem neuromodulation^105^.

Prior work in humans has identified the anterior insula as a key hub that integrates external sensory and internal bodily signals, orchestrating dynamic switching between the default mode network and frontoparietal attention networks^97,106^. Animal studies similarly suggest that noradrenaline release in response to salient, behaviorally relevant events facilitates cortical network reconfiguration^5,107^. Integrating these lines of research, and in correspondence with direct bidirectional projections between the anterior insula and locus coeruleus^9,108^, it has been proposed that the noradrenergic innervation of the anterior insula and related structures facilitates transitioning to network states optimized for new environmental demands^6,107^. Consistent with this framework, our data suggest a functional loop in which stimulus salience is evaluated in frontal areas such as the anterior insula^2^, which recruits noradrenergic neuromodulation to drive network shifts, including insula-mediated default mode network deactivation and dorsal attention network activation^6,97,106,107^. On an electrophysiological level, these network transitions may be coupled with increases in cortical excitability^4^.

Notably, beyond the action-mode network, also the salience network includes hub regions in the anterior insula and cingulate cortex^109^, and in some atlases networks with these names are overlapping^24^. Similar to the action-mode network, salience network activity has been linked to ascending neuromodulatory systems, including noradrenaline^110^. While our study cannot conclusively disentangle the spatially adjacent and functionally linked networks, the more posterior cingulate and more dorsal anterior insula activation favors an action-mode network involvement (^22,109^, see Figure 3).

### Locus coeruleus and action-mode network together support implicit learning

Finally, we examined behavioral adaptations over the course of the experiments. Participants, initially unfamiliar with the task structure, gradually optimized their response times to reflect stimulus probabilities, indicating implicit learning. Optimized response times may arise from preparatory movement initiation, resulting in quicker responses for expected stimuli, but slower responses for unexpected stimuli. Crucially, individuals with greater MRI-indexed locus coeruleus integrity, stronger task-related anterior insula activation, and more pronounced pupil dilation showed greater differentiation in response times between frequent and infrequent stimuli. This suggests that the locus coeruleus and its cortical targets play a key role in learning and adapting to novel environments, matching previous findings from rodents demonstrating that noradrenergic activity is causal for task execution *and* optimization^111^. In addition, our observations are in line with a recent study in humans relating higher MRI-indexed locus coeruleus integrity to larger practice effects^112^ (quicker response times over the course of one year for multiple cognitive tasks), which may be considered a type of uninstructed performance optimization similar to the one observed in this study. Our results moreover support and extend earlier work using *explicit* cognitive assessments to link noradrenergic markers to attention^38,40^, learning, and memory^41,43^.

Please note that what we refer to as implicit learning in this manuscript is how initially naïve participants pick up the statistical regularities of a novel (task) environment without being explicitly instructed^113,114^. Similar concepts have been discussed under the terms *structure learning*^115^ or *statistical learning*^116,117^ in other work. We do not make any assumptions if participants eventually become aware of these regularities^113^. On a neural level, implicit relative to explicit learning is thought to be linked to partially distinct brain responses^113,118–120^ (e.g., in terms of the dependency on medial temporal lobe and striatal circuits, or specific neuromodulators) and, on a cognitive level, differences in the intention to learn^113,120^. The latter may differentially involve neuromodulation-influenced strategic or motivational processes^121^. Finally, while age differences in explicit learning are well established (e.g., in episodic memory^122^), some forms of implicit learning show less pronounced age-related declines^120^.

### Late-life age differences in implicit learning and locus coeruleus-related measures

Post-mortem research identified the locus coeruleus as one of the starting points of Alzheimer’s-related tau accumulation^36^, which may dysregulate noradrenergic activity with increasing age^31,34^ and impair late-life cognition^13,33^. Here we found that a latent measure reflecting locus coeruleus integrity as well as pupil dilation and insula activation to salient stimuli did not differ between younger and older adults. This may indicate that, compared to self-initiated processing, externally-driven noradrenergic responses may be less impaired in aging^38,123,124^. In addition, we found that this noradrenergic latent variable similarly explained implicit learning across age groups, highlighting its behavioral relevance over the adult lifespan. Post-hoc age-group comparisons for each of the variables included in the brain–behavior analyses (cf. Supplementary Information) revealed little differences, apart from slightly larger evoked pupil responses to oddball stimuli in aging.

### Limitations

Small subcortical neuromodulatory nuclei are challenging to assess non-invasively in humans^47,48^. Thus, studies frequently rely on proxy measures, such as pupil size changes that, however, have been linked to multiple neuromodulators^53,54,68^ (also see Figure 4). Adding additional modalities (EEG, MRI, PET), we find correlational evidence for a robust involvement of several brainstem–midbrain neuromodulatory systems (noradrenergic, dopaminergic, serotonergic) that are known to co-activate and interact^3,125^. Future work, optimally at ultra-high field strength and higher spatial resolution, is required to better characterize their crosstalk, potential sex differences therein^126^, and how this regulates behavior.

### Conclusions

By combining neuromodulation-sensitive MRI, concurrent pupillometry–fMRI, and PET-derived neuromodulatory transporter maps, our study suggests the noradrenergic locus coeruleus may contribute to a transition from default-mode to action-mode network activation. This mechanism may support adapting behavior to novel environments, reinforcing the importance of neuromodulatory systems in attention, learning and memory.

## 4. Material and methods

### Study design

Data were collected as part of a larger project investigating neural and cognitive correlates of age-related differences in the noradrenergic system (for details, see^38^). Participants attended three consecutive days of lab-based testing, with the conditioned oddball task administered on the second and third days (Figure 1). On the second day, concurrent pupillometry and electroencephalography (EEG) data were recorded during task performance, while third-day assessments incorporated MRI and pupillometry measurements. To enhance the salience of oddball stimuli, each assessment day included a preceding fear conditioning session, during which the oddball stimuli were repeatedly paired with aversive electrical stimulation. The institutional review board of the German Psychological Association approved the study protocol.

### Participants

The final sample included 39 younger adults (YAs; mean age ± standard deviation: 25.23 ± 3.23 years; range: 20.17–31 years) and 38 older adults (OAs; mean age: 70.61 ± 2.71 years; range: 65.50–75.92 years), all of whom were male. To reduce inter-individual variability, only male participants were included. Pilot testing revealed reliable sex differences in self-selected stimulation intensity during the fear-conditioning procedure, consistent with prior reports of sex-related differences in fear learning and maintenance^127–129^. Such differences may relate in part to variation in the noradrenergic system^126,130^. Restricting the sample to males allowed tighter experimental control, however, future studies should examine whether the present findings generalize across sexes. Two younger adults were excluded from the study after the first assessment day due to low-quality pupil data. Six participants (three per age group) did not participate in the fMRI part of the study (n = 4) or had incomplete eye tracking data (n = 2), leaving a sample of n = 71 for fMRI–pupillometry analyses. All participants were healthy, right-handed, MRI-compatible, fluent German speakers with normal or corrected-to-normal vision, for further characteristics, see^38^. All participants provided written informed consent and were reimbursed for their participation. Exclusion criteria included the use of centrally active drugs, particularly medications affecting the noradrenergic system (e.g., beta-blockers). We did not run formal power calculations before the study was conducted, but based our recruitment on a recent comparable age-comparative study on the noradrenergic system that also used fear conditioning, pupil size and BOLD fMRI measurements^40^.

### Experimental procedures and stimuli

#### Fear conditioning

Participants underwent fear conditioning on each testing day to increase the salience of experimental stimuli and their likelihood of eliciting locus coeruleus activation^40,44–46^. During this phase, a sinusoidal luminance pattern (Gabor patch; CS+) with either horizontal or vertical orientation (0° or 90°) was paired with an aversive electrical shock (US) on 80% of trials. A perceptually matched control stimulus (CS–) with the alternate orientation (vertical or horizontal) was never paired with the shock. Stimulus orientation–reinforcement pairings remained consistent for each participant across days and were counterbalanced within age groups (younger adults [YA]: 21:18; older adults [OA]: 20:18).

Each conditioning session comprised 40 trials, with 20 presentations each of CS+ and CS– in pseudorandomized order across assessment days. Trials began with a one-second fixation cross, followed by a two-second visual stimulus (CS+ or CS–). Shocks were delivered for 0.5 seconds immediately after CS+ offset via a bipolar current stimulator (DS5; Digitimer) connected to a ring electrode affixed to the participant’s left or right index finger. Hand assignment was counterbalanced within age groups (YA: 19:20; OA: 19:19). Each trial concluded with a six-second inter-trial interval, allowing pupil responses to return to baseline.

Participants selected an individually calibrated stimulation intensity deemed unpleasant but not painful before each session^38^. Throughout all conditioning sessions, pupil dilation and gaze position were recorded, and external distractions were minimized.

A procedural error occurred for one older participant, resulting in a reversal of stimulus orientation–shock assignment between the first and second assessment days. This error reduced the likelihood of detecting differences between conditions (CS+/CS–; i.e., worked against our hypothesis), but the participant’s data were retained for analysis to maintain a larger sample size.

### Conditioned oddball task

On the second and third assessment days (EEG–pupillometry and fMRI–pupillometry, respectively), participants completed a modified version of a conditioned oddball task previously shown to increase locus coeruleus activity^44^.

Specifically, on each trial participants viewed a Gabor patch in one of three possible orientations (horizontal, diagonal, vertical; [0°, 45°, 90°]). Two of these orientations (0°, 90°) were extensively familiarized during preceding fear conditioning (see above) and other tasks within the larger project (see^38^), whereas the third orientation (45°) benefited from relative novelty, hypothesized to enhance neuromodulatory activity^131^.

In our experiment, we were interested in visual stimuli that were (1) matched for luminance, (2) easy to discern, (3) inherently neutral and (4) did not produce distracting visual after-effects (testing took place with lights turned off). The Gabor patches satisfied all of these requirements. By a simple tilt (0°, 45°, 90°) of the Gabor patch, we were able to generate three luminance-matched stimulus categories (Standard (CS–), Oddball CS+, Oddball novel). The spatial frequency of the Gabor patches was not linked to any experimental condition and was constant for all stimulus categories.

During the conditioned oddball task, one of the stimulus orientations was presented frequently (standard trials; 0° or 90°), while the two other orientations were rarely presented (oddballs). In total, the task consisted of 160 trials presented in pseudorandomized order that differed over assessment days, with the standard, non-conditioned stimulus (CS–) appearing on 70% of trials (112 trials), while the two oddball stimuli (CS+ and the novel 45° orientation) appeared less frequently, each comprising 15% of trials (24 each).

Gabor patches (0°, 45°, and 90° orientation) were displayed for two seconds followed by an inter-trial interval ranging from 2–12 seconds (mean 3.625 seconds; skewed distribution generated with OptSeq2, http://surfer.nmr.mgh.harvard.edu/optseq/).

During Gabor patch presentation, participants were instructed to respond as quickly and accurately as possible by pressing a button corresponding to the orientation of each stimulus using the hand not used for fear conditioning (see above). Accuracy and response times were recorded for each trial.

### Physiological data recording and preprocessing

#### Eye tracking

During all sessions, participants’ pupil dilation was measured as an indirect marker of central neuromodulatory activity^53–55^ alongside gaze position, using a video-based infrared eye tracker (SR Research EyeLink 1000). On the second day of assessments (during EEG), the desktop mount configuration was used, while on the third day (during fMRI), the long-range mount was employed. The systems operated in a monocular configuration with a spatial resolution of up to 0.25° and a sampling rate of 1000 Hz.

To minimize head movements, participants were seated with their forehead and chin stabilized at a fixed distance of 53.5 cm from the display during the second day. For the MRI session (third day), the eye tracker was positioned within the scanner bore, and data were collected via a mirror affixed to the head coil, which redirected participants’ view to the stimulus display.

Participants were instructed to maintain central fixation throughout the experiments. Before each experiment, the eye tracker was (re)calibrated using a standard 5-point grid. Calibration was considered successful if fixation errors were under 0.5°.

Eye tracking data were preprocessed and missing samples were imputed using published standardized routines^132^. The percentage of imputed samples was logged as an objective quality measure for potential subsequent participant exclusion (proportion of valid eye tracking samples during the fMRI–pupillometry assessment (mean ± standard error): 0.803 ± 0.024).

### Magnetic resonance imaging

MRI data were collected using a 3T Magnetom TIM Trio (Siemens Healthcare) with a 12-channel head coil.

An axial, T1-weighted neuromodulation-sensitive Fast Spin Echo (FSE) sequence was collected to assess markers for locus coeruleus integrity. The following parameters were used^41,133^: acquisition matrix: 440 × 512, 10 slices, voxel size: 0.5 × 0.5 × 2.5 mm, 20 % gap between slices, repetition time (TR): 600 ms, echo time (TE): 11 ms, flip angle (FA): 120 °, acquisition time: 11:48 min. To increase signal to noise ratios in brainstem imaging, the sequence included four online averages and yielded two images, extracted locus coeruleus parameters were averaged to boost stability of estimates^48^.

In addition, a sagittal T1-weighted Magnetization-Prepared Gradient Echo (MPRAGE) sequence with the following parameters was collected to facilitate co-registration to standard space: acquisition matrix: 256 × 256 × 192, voxel size: 1 mm isotropic, repetition time (TR): 2500 ms, echo time (TE): 4.77 ms, inversion time (TI): 1100 ms, flip angle (FA): 10 °, acquisition time: 9:20 min.

Finally, a T2*-weighted whole-brain Echo Planar Imaging (EPI) sequence was used to track blood oxygenation level dependent (BOLD)-changes during the conditioned oddball task. The following imaging parameters were used: acquisition matrix: 72 × 72, 36 slices, 450 images, voxel size: 3 mm isotropic, repetition time (TR): 2000 ms, echo time (TE): 30 ms, flip angle (FA): 80 °, acquisition time: 15 min. The spatial resolution of the fMRI acquisition is comparatively low relative to the size of the locus coeruleus in analyses of human post-mortem tissue^134^, although comparable to previous 3T fMRI studies of the locus coeruleus^135,136^. Accordingly, inferences regarding locus coeruleus-specific activity should be interpreted with caution. To mitigate this limitation, we complemented the fMRI analyses with convergent multimodal measures, including pupil dilation, EEG, dedicated structural MRI, and PET-derived neuromodulatory transporter maps, which together provide additional constraints on the interpretation of locus coeruleus-related effects.

MPRAGE and EPI data were preprocessed using standardized routines as implemented in HeuDiConv^137^, MRIQC^138,139^, and fMRIprep^140^ and as detailed in the supplementary information. Data were transformed to 2 mm isotropic MNI 152 non-linear 2009c asymmetric space and smoothed with an isotropic 8 mm full-width half-maximum (FWHM) kernel for further analyses, following recommendations for smoothing kernels to be at least twice the voxel resolution^141^. A time series of confounds was derived from head-motion estimates, and global signals (within the white matter, the cerebrospinal fluid (CSF) and whole brain mask). These confounds were expanded with the inclusion of temporal derivatives and quadratic terms for each, yielding a total of 36 confounds that were included in within-participant analyses^142,143^.

### Electroencephalography

EEG was recorded during task performance using 61 Ag/AgCl scalp electrodes mounted in an elastic cap (10–10 system; Brain Products GmbH, Germany) and amplified with BrainAmp DC amplifiers. Signals were sampled at 1000 Hz (0.1–250 Hz hardware bandpass) with AFz as ground and referenced online to the right mastoid. The left mastoid was recorded for offline re-referencing. Horizontal and vertical electrooculograms were acquired using three peri-orbital electrodes. Electrode impedances were kept below 5 kΩ during EEG preparation.

EEG data were preprocessed using EEGLAB^144^, including the Eye-EEG toolbox^145^, FieldTrip^146^, and custom MATLAB scripts (for a detailed description, see^38^). Continuous data were demeaned, re-referenced to linked mastoids, downsampled to 500 Hz, and band-pass filtered (0.2–125 Hz, fourth-order Butterworth). Non-stereotyped artifacts were first removed by visual inspection, followed by independent component analysis to identify and reject components reflecting ocular, muscular and cardiac activity. Data were segmented into stimulus-locked epochs (–2500–2500 ms relative to stimulus onset) and remaining artifacts were detected using automated thresholding procedures^147^. Rejected channels were interpolated using spherical splines. After preprocessing, clean epochs were retained for all subsequent analyses.

Event-related potentials were computed from cleaned epochs after baseline correction (–500–0 ms relative to stimulus onset). Time–frequency representations were obtained for each trial and electrode using complex Morlet wavelet convolution (7-cycle wavelets) between 1 and 40 Hz. Power estimates were calculated across the stimulus-locked time window and expressed as condition-specific changes relative to baseline (–500–0 ms relative to stimulus onset).

### Physiological data analyses

#### Pupillometry analysis

Pupil time series were analyzed using the EEGlab^144^, eye-EEG^145^, and FieldTrip^146^ toolboxes. The continuous pupil time series were segmented into trials surrounding presentation of the Gabor patches (for conditioning and oddball experiments). Within-participant trial-level analyses then contrasted pupil responses across experimental conditions after baseline correction (–500–0 ms relative to stimulus onset). That is, for conditioning experiments, CS+ and CS– trials were contrasted, while for the oddball experiments standard and oddball trials were compared (i.e., collapsing the two oddball categories) using sample-wise independent samples *t*-tests. Within-participant analyses were repeated over n_Bootstraps_ = 50 and the resulting *t*-value time series were averaged^38^. Subsequent across-participant analyses tested for the consistency of within-participant effects on a group level (dependent samples *t*-test against zero), while controlling for multiple comparisons using cluster-based random permutation tests^148^. For more details, see^38^.

### Functional magnetic resonance imaging analysis

fMRI data were analyzed using the Statistical Parametric Mapping (SPM) 12 toolbox^149^ to identify brain responses to oddball stimuli. Within-participant event-related analyses included stick functions for each stimulus onset (standards, oddballs) convolved with a canonical hemodynamic response function as well as its temporal and dispersion derivatives^150^. General linear models contrasted BOLD responses to standard and oddball stimuli (aggregating over the two oddball categories to increase power). Subsequent across-participant analyses evaluated the consistency of within-participants effects on a group level with family-wise error correction for multiple comparisons.

Overlap of observed group-level effects with several published functional network atlases (see Figure 3,^23,63,64^) was evaluated using Dice coefficients and permutation tests, as implemented in the Network Correspondence Toolbox^24^. For overlap estimations, statistical maps were thresholded at *t =* 3.

Follow-up within-participant functional connectivity analyses first calculated trial-by-trial activation patterns for each voxel^82^ and next assessed which brain regions’ time series correlated with the locus coeruleus^83^. For this, a locus coeruleus consensus mask was applied as seed region^81^ that reliably segmented the locus coeruleus-related hyperintensity in this study, as shown in the results. That is, our functional connectivity analyses were centered on the region in which we localized the locus coeruleus using dedicated high-resolution structural scans. We did not define an a-priori target for these functional connectivity analyses but instead evaluated each voxel’s correlation pattern with the locus coeruleus (i.e., seed-to-voxel analysis), followed by a stringent control for multiple comparisons.

We detected a reliable association of locus coeruleus and anterior insula time series. To further characterize the temporal properties of this association, we estimated cross-correlations between TR-level locus coeruleus and insula time series on a participant level (using *crosscorr* in *Matlab*;^30^). To this end, we averaged BOLD signals across voxels within the locus coeruleus consensus mask^81^, as well as anterior insula functional-connectivity cluster. Cluster-based random permutation tests probed the consistency of the participant-level cross-correlation estimates on a group level after de-meaning^148^.

Cross-correlation analyses revealed a reliable temporal association between locus coeruleus and anterior insula time series and provided an estimate of their relative lag structure. However, because the hemodynamic response underlying fMRI signals may be regionally heterogeneous, such lag-based measures cannot be straightforwardly interpreted in terms of directional interactions^151^. To address this limitation, we complemented these analyses with Dynamic Causal Modeling (DCM) for fMRI^85,86^, which uses a generative model of neuronal dynamics and hemodynamic responses to infer directed effective connectivity. By explicitly modeling the latent neuronal processes underlying the observed BOLD signals, DCM allows us to test whether the observed temporal associations reflect directed influences (e.g., insula→locus coeruleus or locus coeruleus→insula) and whether these influences are modulated by task events. Thus, the DCM results provide an extension of the cross-correlation findings.

We modeled directed interactions between two a-priori defined regions of interest: the anterior insula and the locus coeruleus, using the same anatomical masks as in the main connectivity analyses. For each participant, regional time series were extracted in SPM12 as the first principal eigenvariate of voxels showing task-related responses (*F*-contrast including standard and oddball regressors convolved with the canonical hemodynamic response function and its temporal derivatives), while controlling for nuisance regressors. Deterministic task-based DCM was implemented following current recommendations^85,86^.

Each participant’s model contained two nodes (anterior insula and locus coeruleus) with reciprocal intrinsic connections and region-specific self-connections (A matrix). Stimulus onsets served as driving inputs to both regions (C matrix), and the oddball condition was allowed to modulate regional self-connections, indexing task-related changes in local excitability (B matrix). Inputs were not mean-centered for the primary analyses, but were for sensitivity analyses.

Models were estimated using Variational Laplace. Group inference on connectivity parameters (A and B matrix parameters) was performed using Parametric Empirical Bayes (PEB), which iteratively re-estimates individual models using group-level priors. Bayesian Model Reduction (BMR) with greedy search identified the most parsimonious model, and parameter estimates were obtained using Bayesian Model Averaging (BMA). Posterior probabilities (Pp) > 0.95 were interpreted as strong evidence and Pp > 0.75 as positive evidence.

To assess robustness for a small brainstem structure, DCM was estimated using both default and reduced spatial smoothing data (8 mm and 2 mm FWHM).

### Structural magnetic resonance imaging analysis

Neuromodulation-sensitive FSE images were analyzed using a previously established semi-automatic procedure to extract proxies for locus coeruleus integrity^41,81,133^ using Advanced Normalization Tools (version 2.3.3.; ANTs^152,153^).

In brief, first MPRAGE scans were linearly resampled to 0.5 mm isotropic resolution before estimating a whole-brain group template (six iterations; *antsMultivariateTemplateConstruction,* including *N4BiasFieldCorrection*). Next, native-space FSE scans were non-linearly aligned to within-person template-space MPRAGE images using *antsRegistrationSyNQuick* to facilitate subsequent brainstem template construction based on the outputs of this transformation (six iterations; *antsMultivariateTemplateConstruction,* including *N4BiasFieldCorrection*). Finally, the brainstem template was moved to 0.5 mm linear MNI space using the whole-brain template as an intermediate target (*antsRegistration,* facilitated by a custom coregistration mask). All transformation matrices were then concatenated and applied to individual neuromodulation-sensitive FSE scans, bringing them directly from native to MNI space in a single step (*antsApplyTransforms*).

To extract proxies for locus coeruleus integrity, standard space FSE images were masked using a high-confidence locus coeruleus consensus mask^81^ that captured the observed locus coeruleus hyperintensity well (see Figure 7). To normalize intensity values for across-participant analyses, scans were also masked with a pontine reference area mask^81^. The peak voxel intensity for each slice of each region of interest was then automatically identified. Finally, intensity ratios for each slice of the locus coeruleus were computed as a measure of structural integrity (see Equation 1). For each slice, these ratios were derived by dividing the difference between the peak intensity in the locus coeruleus and the corresponding reference region by the peak intensity of the reference region, following previously established _methods41,81,133._

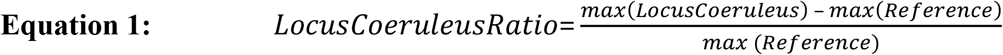

For subsequent analyses, extracted slice-wise values were averaged over hemispheres and then the peak value across slices was selected, while excluding the two most rostral slices to avoid unreliable intensity values at the edge of the acquisition^41^.

### Electrophysiological analysis

Statistical analyses of EEG data followed the same general framework as described above for pupillometry. Segmented baseline-corrected EEG signals were contrasted between experimental conditions (oddball vs. standard stimuli) separately for event-related potentials and time–frequency power estimates. For each participant, condition differences were computed sample-wise using independent-samples *t*-tests, yielding spatiotemporal (electrode × time) maps for event-related potential data and spectro-spatiotemporal (electrode × frequency × time) maps for time–frequency power data. To minimize biases due to unequal trial counts, contrasts were repeated across n_Bootstraps_ = 50 random subsets of trials and the resulting *t*-value maps were averaged.

Group-level inference assessed the consistency of within-participant effects using dependent-samples *t*-tests against zero. Multiple comparisons across electrodes, time and frequency were controlled using non-parametric cluster-based permutation testing as implemented in FieldTrip. The procedure was identical for event-related potential and time–frequency measures, differing only in dimensionality (two- vs. three-dimensional data).

### Behavioral data analysis

In the conditioned oddball task, participants were instructed to press a specific button corresponding to the orientation of the presented Gabor stimulus. The majority of trials (70%) featured the standard stimulus, creating a high probability that a standard stimulus would follow another standard stimulus. We assumed that due to this consistency participants would build up a tendency (bias) to respond with the corresponding key. We termed these *stay-standard* trials. Occasionally, this expectation was violated by the presentation of one of the oddball stimuli, requiring participants to switch response keys (*switch-to-oddball trials*). Following an oddball stimulus, most trials returned to the standard stimulus (*switch-to-standard*), while less frequently, another oddball appeared (*stay-oddball*). Following this reasoning, we estimated mean response times for each of these four trial categories for each participant (2 (stay/switch) × 2 (standard/oddball)) after log-transformation and *Z*-standardization of response times across trials.

Crucially, participants began the task without prior knowledge of the statistical structure or trial distribution. The extent to which participants adjusted their behavior over time in response to the predictability of the stimuli is thus interpreted as an implicit measure of (task structure) learning. To quantify if participant indeed learned the statistical structure of the task, we down-sampled the trial-level response -time time series of the repeated oddball experiments (320 trials in total, two repetitions of 160 trials each; cf. Figure 8). That is, we averaged response times over 10 adjacent trials within the four trial categories, yielding 32 response-time time bins, which we contrasted across trial categories. Finally, we evaluated if participants over the course of the experiment developed a tendency to preferentially focus on one trial category at the expense of another. To this end, we probed if the contrast of response times (*switch-to-oddball* – *stay-standard*) increased over time. We repeated these analyses on a group level (correlation of group-averaged response time contrast and time bin) and on a single-participant level (linear regression predicting response time contrasts by time bin).

Finally, participant-level beta estimates (quantifying implicit learning) were contrasted against zero using dependent**-**samples *t*-test to evaluate the consistency of implicit learning on a group level.

### Multimodal data analyses

#### Combined pupillometry–functional magnetic resonance imaging analyses

Presentation of oddball stimuli elicited pupil dilation and distributed cortical activation, including the action-mode network. To test which brain regions’ activation scaled with pupil-indexed neuromodulation, we estimated a separate set of general linear models. First, we down-sampled each participant’s pupil time series to the fMRI temporal resolution (TR = 2 s) and added it as a regressor to the participant-level models^154^. The same confounds as in unimodal fMRI analyses were applied (see above), while task regressors were omitted. Subsequent analyses across participants assessed the consistency of within-participant effects at the group level, applying family-wise error correction to account for multiple comparisons. Subsets of these analyses were restricted to participants with high-quality pupil data (>50% valid non-imputed samples; 64/71 participants^132^).

#### Association of pupillometry–functional magnetic resonance imaging patterns and PET-derived transporter maps

Combined pupillometry–fMRI analyses showed that brainstem–midbrain and insula activation correlated with pupil size variations, suggesting neuromodulatory influences. To more directly test this, we leveraged publicly available maps of neuromodulatory transporter distributions^56^. These PET maps were not collected from the participants in our sample.

In line with the smooth, low-resolution nature of these PET maps (cf. Figure 5 a/c), we did not intend to explore the transporter distribution in small neuromodulatory nuclei. Instead, after observing a reliable coupling between moment-to-moment pupil fluctuations and BOLD activity (cf. Figure 4), pointing to a neuromodulatory contribution, we wanted to clarify which neuromodulatory systems may give rise to this *brain-wide* activity pattern. This analysis is in part motivated by the known association of pupil size with multiple neuromodulatory systems^54,65,66,68^. That is, we here focused primarily on terminal regions of neuromodulatory systems (e.g., the action-mode network, cf. Figure 4), in which, after release, neuromodulators are transported back into the presynaptic cell, testing if the voxel-wise magnitude of pupil-linked BOLD activation was linked to the transporter expression of various neuromodulatory systems. The underlying assumption is that if the brain-wide activity pattern depends on a certain neuromodulatory system, it should spatially overlap with the distribution of its transporters, which give rise to it.

All PET-derived maps and our pupil–fMRI activation maps were resampled to a matching resolution and common space (1 mm, MNI-ICBM 152 non-linear 2009 version c space). Next, we probed the voxel-wise association of noradrenergic transporter expression and pupil-linked BOLD activation using two alternative transporter maps using Pearson’s correlations^56,70–74^. To judge the reliability of these effects, we ran two negative control analyses, using two maps of GABAergic receptor distribution acquired from two different sources^75,76^ as well as two comparative analyses, using maps of the dopaminergic ^78^ and serotonergic transporter^56^ distribution. For each of these, observed effects were contrasted to reference distributions of correlation coefficients estimated with permuted pupillometry-fMRI data while preserving the spatial autocorrelation of the observed data. Specifically, we applied a phase-randomization approach, which retains the power spectrum of the original data while randomizing its phase in the Fourier domain^155^. This method generates surrogate datasets that maintain the autocorrelation properties of the original data (i.e., the pupil–fMRI activation map) while breaking voxel-wise correspondences with the PET-derived transporter/receptor maps. We computed correlation coefficients between the shuffled pupil-linked BOLD activation pattern and the transporter/receptor maps across 10,000 iterations, forming a null distribution, to which we compared the observed correlation coefficients.

Finally, to determine if the voxel-wise noradrenergic transporter expression explained variance in pupil-linked BOLD patterns (*PupilBOLD*), we used multiple regression analyses (using *fitglm* in Matlab). Specifically, we compared a base model (*PupilBOLD* ∼ 1 + *5HTT* + *DAT*) incorporating the serotonergic (*5HTT*) and dopaminergic (*DAT*) transporter expression to a full model including all three neuromodulatory systems (*PupilBOLD* ∼ 1 + *5HTT* + *DAT* + *NAT*). To obtain a single noradrenergic transporter predictor (*NAT*), the two available maps were averaged^70,74^. Differences in model fit between base and full models were evaluated using a likelihood-ratio test and compared to a null distribution obtained from adding shuffled noradrenergic transporter predictors (10,000 iterations), yielding corrected *p*-values.

In addition to voxel-wise spatial correlations, we quantified overlap using Dice coefficients. Pupil–BOLD and PET maps were first standardized (*z*-scored) and then thresholded (z ≥ 1.96). To facilitate a comparable standardization, all PET maps were first rescaled to a value range of 0–1 and voxels falling below 0.05 were removed from subsequent analyses. Then the spatial overlap between binarized maps was computed. Statistical significance was determined by comparing observed Dice values to phase-randomization reference distributions generated as described above. To evaluate neuromodulatory system-specific contributions where transporter maps exhibited spatial overlap, partial Dice coefficients were computed. That is, for a given transmitter map, voxels exceeding threshold in other transporter maps were excluded prior to overlap estimation, thereby isolating unique spatial correspondence. For comparability across maps with differing spatial autocorrelation properties, corrected correspondence metrics were additionally reported by normalizing observed Dice coefficients relative to the mean of the corresponding null distribution.

### Combined pupillometry–EEG analyses

Associations between pupil dilation and EEG responses were evaluated within participants on a trial-by-trial basis. For each spatiotemporal EEG sample, correlation coefficients relating averaged and standardized pupil estimates to event-related potential or time–frequency power were computed, resulting in participant-specific correlation maps. Group-level significance was assessed using the same cluster-based permutation framework described in *Pupillometry analysis*, testing correlation values against zero. This approach isolates neuromodulation-linked fluctuations shared across modalities. For more details, see_38._

### Combined EEG–MRI analyses

To test whether the identified electrophysiological and hemodynamic signatures reflected a shared neuromodulatory process, we related pupil–EEG and pupil–fMRI coupling across participants. Pupil measurements were obtained in both sessions and served as a physiological bridge between modalities (see Figure 1–2). Low pupil quality participants were excluded from these analyses (see above).

For the EEG data, participant-specific statistical maps quantifying trial-by-trial associations between pupil size and time–frequency power (electrode × frequency × time) were first computed as described above. To obtain a single summary measure per participant, the high-dimensional pupil–EEG coupling data were vectorized and reduced using principal component analysis (PCA). The first principal component score was retained as an index of pupil-linked cortical excitability modulation strength.

For the fMRI data, each participant’s pupil-linked BOLD response was represented by voxel-wise regression coefficients from the pupil–fMRI general linear model (see above), with CSF regions masked out. These maps were reshaped into participant × voxel matrices and related to the EEG summary index using partial least squares correlation (cf. *Combined brain–behavior analyses*). Statistical significance of latent variables was assessed using permutation testing, and the reliability of voxel contributions was estimated using bootstrap resampling (reported as bootstrap ratios; BSR).

To aid interpretation, spatial correspondence between the resulting bootstrap ratio maps, expressing the voxel-wise loadings to the latent EEG–fMRI association, and canonical functional networks was quantified using Dice overlap coefficients with established network parcellations^24^. Because EEG and fMRI were recorded in separate sessions, the analysis was performed at the participant level rather than trial level. Thus, the reported associations reflect individual differences in neuromodulation-linked coupling shared across modalities.

### Combined brain–behavior analyses including pupillometry and MRI

Our physiological analyses yielded multiple locus coeruleus-related measures (MRI-indexed locus coeruleus integrity; pupil-indexed neuromodulation; locus coeruleus-linked action-mode network activation). Given the prominent involvement of the noradrenergic system in learning and memory ^3^, we next tested if these explained individual differences in implicit learning during the conditioned oddball task, using partial least squares correlations _(PLSC156,157)._

In short, partial least squares correlation is a multivariate statistical technique used to identify patterns of covariance between two sets of variables. In this study, behavioral partial least squares correlation was applied to explore the relationship between locus coeruleus-related measures and task-derived measures of implicit learning. By decomposing the covariance structure with Singular Value Decomposition, partial least squares correlation identifies latent variables (LVs) that maximize the shared variance between the datasets (i.e., locus coeruleus-related measures and implicit learning). Each latent variable is characterized by a pair of weighted combinations of variables—one from each dataset—allowing for the assessment of how strongly these patterns are expressed across participants. Statistical significance of the latent variables was determined using permutation tests (n_permutations_ = 10,000), and the reliability of individual variable contributions was evaluated with bootstrapping (n_bootstraps_ = 10,000). A ratio of a variable’s weight and its bootstrapped standard error, termed bootstrap ratio, (BSR), allows identifying indicators reliably contributing to the latent variable (|BSR| ≥ 1.96, interpreted analogous to *z*-scores). Taken together, partial least squares correlation provides a robust method for examining complex, high-dimensional relationships, integrating locus coeruleus-related and learning-related measures. For further details, see^81^.

The following neural and behavioral variables were used as input to the partial least-squares analyses: (1) individual task-related left anterior insula activations (for Oddball > Standard trials), extracted using MarsBar^158^, averaged over voxels of the cluster that showed reliable associations with pupil-linked neuromodulation (Figure 4); (2) individual task-related pupil dilation (for Oddball > Standard trials), extracted using the cluster shown in Figure 2 and averaged over samples; (3) peak locus coeruleus contrast ratios, averaged over hemispheres; and (4–8) individual response times for each trial category, 2 (stay/switch) × 2 (standard/oddball), averaged over trials after log-transformation and *z*-standardization, as shown in Figure 8. All input variables were standardized across individuals, and participants with absolute values >3 were dropped before analyses (n = 3). Post-hoc independent samples *t*-tests probed age differences in the latent brain and behavioral variables. As a sensitivity analysis, we replicated partial least squares correlation findings using a simpler analytical strategy that averaged over standardized neural indicators (i.e., calculated composite scores based on (1)-(3), see PLSC analyses), estimated the difference in response times for oddball and standard trials, and then correlated these measures.

### Combined brain–behavior analyses including pupillometry and EEG

To also assess the behavioral relevance of cortical excitability adjustments, we related pupil-linked EEG responses to implicit learning performance. Using the same dimensionality-reduction approach as in the cross-modal analyses (see *Combined EEG–MRI analyses*), participant-specific pupil–EEG coupling maps were summarized by the first principal component, yielding a single index of pupil-linked cortical excitability modulation per participant. Implicit learning was quantified as response-time differences between oddball and standard trials, reflecting sensitivity to the task’s statistical structure. Pearson correlations were then computed to test whether individual differences in pupil-linked excitability modulation were associated with behavioral adaptation.

## Supporting information

Supplementary methods and results

## Acknowledgements

MW-B received support from the German Research Foundation (DFG, WE 4269/5-1) and the Jacobs Foundation (Early Career Research Fellowship 2017–2019).

MM’s work was supported by an Alexander von Humboldt fellowship and by National Institutes of Health grant R01AG025340 and R01AG080652.

During the work on this article, MJD was a member of the International Max Planck Research School on Computational Methods in Psychiatry and Ageing Research (IMPRS COMP2PSYCH, https://www.mps-uclcentre.mpg.de/comp2psych. Participating institutions: Max Planck Institute for Human Development, University College London). At the time of writing, TL was a PhD student in the Max Planck School of Cognition.

Acknowledgment is made to the donors of the ADR A2024006F, a program of the BrightFocus Foundation, for support of this research (MJD).

This work was in part conducted at the Max Planck Dahlem Campus of Cognition (MPDCC) of the Max Planck Institute for Human Development, Berlin, Germany.

## Author contributions

MJD, MM and MWB designed the study. MJD performed the experiments together with a team of research assistants. MJD analyzed the data. TL contributed to a subset of analyses. MJD wrote the paper. All authors gave conceptual advice and revised the paper.

## Data availability

Participants in this study did not consent to the public sharing of their data. To obtain access to participant-level data, please contact the corresponding author (MJD) with a signed data sharing agreement (template available at: https://osf.io/4t5hj/files/osfstorage). Un/thresholded group-level data are available at https://osf.io/4t5hj/.

## Material and correspondence

Correspondence and requests for materials should be addressed to Martin J. Dahl.

## References

1. Uddin, L. Q. Cognitive and behavioural flexibility: neural mechanisms and clinical considerations. Nat. Rev. Neurosci. 1–13 (2021) doi:10.1038/s41583-021-00428-w.

2. Mather, M., Clewett, D., Sakaki, M. & Harley, C. W. Norepinephrine ignites local hotspots of neuronal excitation: How arousal amplifies selectivity in perception and memory. Behavioral and Brain Sciences 39, e200 (2016).

3. Sara, S. J. The locus coeruleus and noradrenergic modulation of cognition. Nat. Rev. Neurosci. 10, 211–223 (2009).

4. Dahl, M. J., Mather, M. & Werkle-Bergner, M. Noradrenergic modulation of rhythmic neural activity shapes selective attention. Trends Cogn. Sci. 26, 38–52 (2022).

5. Bouret, S. & Sara, S. J. Network reset: A simplified overarching theory of locus coeruleus noradrenaline function. Trends Neurosci. 28, 574–582 (2005).

6. Corbetta, M., Patel, G. & Shulman, G. L. The reorienting system of the human brain: From environment to theory of mind. Neuron 58, 306–324 (2008).

7. Dayan, P. & Yu, A. J. Phasic norepinephrine: A neural interrupt signal for unexpected events. Network: Computation in Neural Systems 17, 335–350 (2006).

8. Nassar, M. R. Toward a computational role for locus coeruleus/norepinephrine arousal systems. Curr. Opin. Behav. Sci. 59, 101407 (2024).

9. Aston-Jones, G. & Cohen, J. D. An integrative theory of locus coeruleus-norepinephrine function: Adaptive gain and optimal performance. Annu. Rev. Neurosci. 28, 403–450 (2005).

10. Thiele, A. & Bellgrove, M. A. Neuromodulation of attention. Neuron 97, 769–785 (2018).

11. Servan-Schreiber, D., Printz, H. & Cohen, J. D. A network model of catecholamine effects: Gain, signal-to-noise ratio, and behavior. Science (1979). 249, 892–5 (1990).

12. Ghosh, S. & Maunsell, J. H. R. Locus coeruleus norepinephrine contributes to visual-spatial attention by selectively enhancing perceptual sensitivity. Neuron 112, 2231–2240.e5 (2024).

13. Dahl, M. J., Kulesza, A., Werkle-Bergner, M. & Mather, M. Declining locus coeruleus-dopaminergic and noradrenergic modulation of long-term memory in aging and Alzheimer’s disease. Neurosci. Biobehav. Rev. 105358 (2023) doi:10.1016/J.NEUBIOREV.2023.105358.

14. Hagena, H., Hansen, N. & Manahan-Vaughan, D. β-adrenergic control of hippocampal function: Subserving the choreography of synaptic information storage and memory. Cerebral Cortex 26, 1349–1364 (2016).

15. Eschenko, O. The role of the locus coeruleus in cellular and systems memory consolidation. in Handbook of behavioral neuroscience: Handbook of in vivo neural plasticity techniques (ed. Manahan-Vaughan, D.) vol. 28 327–347 (Elsevier, London, 2019).

16. Hagena, H. & Manahan-Vaughan, D. Oppositional and competitive instigation of hippocampal synaptic plasticity by the VTA and locus coeruleus. Proceedings of the National Academy of Sciences 122, e2402356122 (2024).

17. Jordan, R. The locus coeruleus as a global model failure system. Trends Neurosci. 47, 92–105 (2024).

18. Jepma, M. et al. Catecholaminergic Regulation of Learning Rate in a Dynamic Environment. PLoS Comput. Biol. 12, e1005171 (2016).

19. Silvetti, M., Vassena, E., Abrahamse, E. & Verguts, T. Dorsal anterior cingulate-brainstem ensemble as a reinforcement meta-learner. PLoS Comput. Biol. 14, e1006370 (2018).

20. Li, T., Marble, H., Chen, T., Razmi, N. & Nassar, M. R. Fluctuations in arousal reflect latent state transitions that facilitate behavioral optimization. bioRxiv 2025.02.06.636887 (2025) doi:10.1101/2025.02.06.636887.

21. Nassar, M. R., Wilson, R. C., Heasly, B. & Gold, J. I. An Approximately Bayesian Delta-Rule Model Explains the Dynamics of Belief Updating in a Changing Environment. Journal of Neuroscience 30, 12366–12378 (2010).

22. Dosenbach, N. U. F., Raichle, M. E. & Gordon, E. M. The brain’s action-mode network. Nature Reviews Neuroscience 2025 1–11 (2025) doi:10.1038/s41583-024-00895-x.

23. Yeo, B. T. T. et al. The organization of the human cerebral cortex estimated by intrinsic functional connectivity. J. Neurophysiol. 106, 1125–1165 (2011).

24. Kong, R. et al. A network correspondence toolbox for quantitative evaluation of novel neuroimaging results. Nature Communications 2025 16:1 16, 1–16 (2025).

25. Dosenbach, N. U. F., Fair, D. A., Cohen, A. L., Schlaggar, B. L. & Petersen, S. E. A dual-networks architecture of top-down control. Trends Cogn. Sci. 12, 99–105 (2008).

26. Berridge, C. W. Noradrenergic modulation of arousal. Brain Res. Rev. 58, 1–17 (2008).

27. Carter, M. E. et al. Tuning arousal with optogenetic modulation of locus coeruleus neurons. Nat. Neurosci. 13, 1526–1535 (2010).

28. Hayat, H. et al. Locus coeruleus norepinephrine activity mediates sensory-evoked awakenings from sleep. Sci. Adv. 6, eaaz4232 (2020).

29. Osorio-Forero, A. et al. Infraslow noradrenergic locus coeruleus activity fluctuations are gatekeepers of the NREM–REM sleep cycle. Nat. Neurosci. https://doi.org/110.1038/s41593-024-01822-0 (2024) doi:10.1038/s41593-024-01822-0.

30. Kucyi, A. & Parvizi, J. Pupillary dynamics link spontaneous and task-evoked activations recorded directly from human insula. Journal of Neuroscience 40, 6207–6218 (2020).

31. Kelberman, M. A. et al. Diversity of ancestral brainstem noradrenergic neurons across species and multiple biological factors. bioRxiv https://doi.org/10.1101/2024.10.14.618224 (2024) doi:10.1101/2024.10.14.618224.

32. Theofilas, P. et al. Locus coeruleus volume and cell population changes during Alzheimer’s disease progression: A stereological study in human postmortem brains with potential implication for early-stage biomarker discovery. Alzheimer’s and Dementia 13, 236–246 (2017).

33. Mooraj, Z. et al. Toward a functional future for the cognitive neuroscience of human aging. Neuron 113, 154–183 (2025).

34. Weinshenker, D. Long road to ruin: Noradrenergic dysfunction in neurodegenerative disease. Trends Neurosci. 41, 211–223 (2018).

35. Mather, M. & Harley, C. W. The locus coeruleus: Essential for maintaining cognitive function and the aging brain. Trends Cogn. Sci. 20, 214–226 (2016).

36. Braak, H., Thal, D. R., Ghebremedhin, E. & Del Tredici, K. Stages of the pathologic process in Alzheimer disease: Age categories from 1 to 100 years. J. Neuropathol. Exp. Neurol. 70, 960–969 (2011).

37. Ehrenberg, A. J. et al. Priorities for research on neuromodulatory subcortical systems in Alzheimer’s disease: Position paper from the NSS PIA of ISTAART. Alzheimer’s & Dementia 8, 21 (2023).

38. Dahl, M. J., Mather, M., Sander, M. C. & Werkle-Bergner, M. Noradrenergic responsiveness supports selective attention across the adult lifespan. Journal of Neuroscience 40, 4372–4390 (2020).

39. Kosciessa, J. Q., Mayr, U., Lindenberger, U. & Garrett, D. D. Broadscale dampening of uncertainty adjustment in the aging brain. Nature Communications 2024 15:1 15, 1–18 (2024).

40. Lee, T.-H. et al. Arousal increases neural gain via the locus coeruleus–noradrenaline system in younger adults but not in older adults. Nat. Hum. Behav. 2, 356–366 (2018).

41. Dahl, M. J. et al. The integrity of dopaminergic and noradrenergic brain regions is associated with different aspects of late-life memory performance. Nature Aging 2023 1–16 (2023) doi:10.1038/s43587-023-00469-z.

42. Bueichekú, E. et al. Spatiotemporal patterns of locus coeruleus integrity predict cortical tau and cognition. Nature Aging 2024 1–13 (2024) doi:10.1038/s43587-024-00626-y.

43. Jacobs, H. I. L. et al. In vivo and neuropathology data support locus coeruleus integrity as indicator of Alzheimer’s disease pathology and cognitive decline. Sci. Transl. Med. 13, eabj2511 (2021).

44. Aston-Jones, G., Rajkowski, J., Kubiak, P. & Alexinsky, T. Locus coeruleus neurons in monkey are selectively activated by attended cues in a vigilance task. Journal of Neuroscience 14, 4467–4480 (1994).

45. Deitcher, Y., Leibner, Y., Kutzkel, S., Zylbermann, N. & London, M. Nonlinear relationship between multimodal adrenergic responses and local dendritic activity in primary sensory cortices. bioRxiv 814657 (2019) doi:10.1101/814657.

46. Wilmot, J. H. et al. Phasic locus coeruleus activity enhances trace fear conditioning by increasing dopamine release in the hippocampus. Elife 12, (2023).

47. Astafiev, S. V., Snyder, A. Z., Shulman, G. L. & Corbetta, M. Comment on ‘Modafinil shifts human locus coeruleus to low-tonic, high-phasic activity during functional MRI’ and ‘Homeostatic sleep pressure and responses to sustained attention in the suprachiasmatic area’. Science (1979). 328, 309 (2010).

48. Forstmann, B. U., De Hollander, G., Van Maanen, L., Alkemade, A. & Keuken, M. C. Towards a mechanistic understanding of the human subcortex. Nat. Rev. Neurosci. 18, 57–65 (2016).

49. Betts, M. J. et al. Locus coeruleus imaging as a biomarker for noradrenergic dysfunction in neurodegenerative diseases. Brain 142, 2558–2571 (2019).

50. Pérot, J.-B. et al. Longitudinal neuromelanin changes in prodromal and early Parkinson’s disease in humans and rat model. bioRxiv 2024.10.22.619619 (2024) doi:10.1101/2024.10.22.619619.

51. Watanabe, T., Tan, Z., Wang, X., Martinez-Hernandez, A. & Frahm, J. Magnetic resonance imaging of noradrenergic neurons. Brain Struct. Funct. 224, 1609–1625 (2019).

52. Keren, N. I. et al. Histologic validation of locus coeruleus MRI contrast in post-mortem tissue. Neuroimage 113, 235–245 (2015).

53. Joshi, S. & Gold, J. I. Pupil size as a window on neural substrates of cognition. Trends Cogn. Sci. 24, 466–480 (2020).

54. Grujic, N., Polania, R. & Burdakov, D. Neurobehavioral meaning of pupil size. Neuron 0, (2024).

55. Privitera, M. et al. A complete pupillometry toolbox for real-time monitoring of locus coeruleus activity in rodents. Nat. Protoc. 15, 2301–2320 (2020).

56. Hansen, J. Y. et al. Mapping neurotransmitter systems to the structural and functional organization of the human neocortex. Nature Neuroscience 2022 1–13 (2022) doi:10.1038/s41593-022-01186-3.

57. Vazey, E. M., Moorman, D. E. & Aston-Jones, G. Phasic locus coeruleus activity regulates cortical encoding of salience information. Proc. Natl. Acad. Sci. U. S. A. 115, E9439–E9448 (2018).

58. Nieuwenhuis, S., Aston-Jones, G. & Cohen, J. D. Decision making, the P3, and the locus coeruleus-norepinephrine system. Psychol. Bull. 131, 510–532 (2005).

59. Neves, R. M., van Keulen, S., Yang, M., Logothetis, N. K. & Eschenko, O. Locus coeruleus phasic discharge is essential for stimulus-induced gamma oscillations in the prefrontal cortex. J. Neurophysiol. 119, 904–920 (2018).

60. Weijs, M. L. et al. Modulating cortical excitability and cortical arousal by pupil self-regulation. Nature Communications 2025 16:1 16, 1–19 (2025).

61. McCormick, D. A. Cholinergic and noradrenergic modulation of thalamocortical processing. Trends Neurosci. 12, 215–221 (1989).

62. George, M. S. & Aston-Jones, G. Noninvasive techniques for probing neurocircuitry and treating illness: vagus nerve stimulation (VNS), transcranial magnetic stimulation (TMS) and transcranial direct current stimulation (tDCS). Neuropsychopharmacology 2010 35:1 35, 301–316 (2009).

63. Gordon, E. M. et al. Precision Functional Mapping of Individual Human Brains. Neuron 95, 791–807.e7 (2017).

64. Ji, J. L. et al. Mapping the human brain’s cortical-subcortical functional network organization. Neuroimage 185, 35–57 (2019).

65. Cazettes, F., Reato, D., Morais, J. P., Renart, A. & Mainen, Z. F. Phasic activation of dorsal raphe serotonergic neurons increases pupil size. Current Biology 31, 192–197.e4 (2020).

66. Reimer, J. et al. Pupil fluctuations track rapid changes in adrenergic and cholinergic activity in cortex. Nat. Commun. 7, 13289 (2016).

67. Joshi, S., Li, Y., Kalwani, R. M. & Gold, J. I. Relationships between pupil diameter and neuronal activity in the locus coeruleus, colliculi, and cingulate cortex. Neuron 89, 221–234 (2016).

68. Meissner, S. N. et al. Self-regulating arousal via pupil-based biofeedback. Nature Human Behaviour 2023 1–20 (2023) doi:10.1038/s41562-023-01729-z.

69. de Gee, J. W. et al. Dynamic modulation of decision biases by brainstem arousal systems. Elife 6, 1–36 (2017).

70. Hesse, S. et al. Central noradrenaline transporter availability in highly obese, non-depressed individuals. Eur. J. Nucl. Med. Mol. Imaging 44, 1056–1064 (2017).

71. Belfort-Deaguiar, R. et al. Noradrenergic Activity in the Human Brain: A Mechanism Supporting the Defense Against Hypoglycemia. J. Clin. Endocrinol. Metab. 103, 2244–2252 (2018).

72. Sanchez-Rangel, E. et al. Norepinephrine transporter availability in brown fat is reduced in obesity: a human PET study with [11C] MRB. International Journal of Obesity 2019 44:4 44, 964–967 (2019).

73. Li, C. R. et al. Decreased norepinephrine transporter availability in obesity: Positron Emission Tomography imaging with (S,S)-[11C]O-methylreboxetine. Neuroimage 86, 306–310 (2014).

74. Ding, Y. S. et al. PET imaging of the effects of age and cocaine on the norepinephrine transporter in the human brain using (S,S)-[(11)C]O-methylreboxetine and HRRT. Synapse 64, 30–38 (2010).

75. Dukart, J. et al. Cerebral blood flow predicts differential neurotransmitter activity. Scientific Reports 2018 8:1 8, 1–11 (2018).

76. Nørgaard, M. et al. A high-resolution in vivo atlas of the human brain’s benzodiazepine binding site of GABAA receptors. Neuroimage 232, (2021).

77. Breton-Provencher, V. & Sur, M. Active control of arousal by a locus coeruleus GABAergic circuit. Nat. Neurosci. 22, 218–228 (2019).

78. Sasaki, T. et al. Quantification of dopamine transporter in human brain using PET with 18F-FE-PE2I. J. Nucl. Med. 53, 1065–1073 (2012).

79. Hesse, S. et al. Central noradrenaline transporter availability in highly obese, non-depressed individuals. Eur. J. Nucl. Med. Mol. Imaging 44, 1056–1064 (2017).

80. Ding, Y. S. et al. PET imaging of the effects of age and cocaine on the norepinephrine transporter in the human brain using (S,S)-[(11)C]O-methylreboxetine and HRRT. Synapse 64, 30–38 (2010).

81. Dahl, M. J. et al. Locus coeruleus integrity is related to tau burden and memory loss in autosomal-dominant Alzheimer’s disease. Neurobiol. Aging 112, 39–54 (2022).

82. Mumford, J. A., Turner, B. O., Ashby, F. G. & Poldrack, R. A. Deconvolving BOLD activation in event-related designs for multivoxel pattern classification analyses. Neuroimage 59, 2636–2643 (2012).

83. Rissman, J., Gazzaley, A. & D’Esposito, M. Measuring functional connectivity during distinct stages of a cognitive task. Neuroimage 23, 752–763 (2004).

84. Box, G. E. P., Jenkins, G. M. & Reinsel, G. C. Time Series Analysis: Forecasting and Control. (Prentice Hall, Englewood Cliffs, N.J., 1994).

85. Zeidman, P. et al. A guide to group effective connectivity analysis, part 1: First level analysis with DCM for fMRI. Neuroimage 200, 174–190 (2019).

86. Zeidman, P. et al. A guide to group effective connectivity analysis, part 2: Second level analysis with PEB. Neuroimage 200, 12–25 (2019).

87. Krishnan, A., Williams, L. J., McIntosh, A. R. & Abdi, H. Partial Least Squares (PLS) methods for neuroimaging: A tutorial and review. Neuroimage 56, 455–475 (2011).

88. McGinley, M. J. et al. Waking State: Rapid Variations Modulate Neural and Behavioral Responses. Neuron vol. 87 1143–1161 Preprint at 10.1016/j.neuron.2015.09.012 (2015).

89. Stitt, I., Zhou, Z. C., Radtke-Schuller, S. & Fröhlich, F. Arousal dependent modulation of thalamo-cortical functional interaction. Nat. Commun. 9, 2455 (2018).

90. Nestvogel, D. B. & McCormick, D. A. Visual thalamocortical mechanisms of waking state-dependent activity and alpha oscillations. Neuron https://doi.org/10.1016/J.NEURON.2021.10.005 (2021) doi:10.1016/J.NEURON.2021.10.005.

91. Bestmann, S., Ruff, C. C., Blakemore, C., Driver, J. & Thilo, K. V. Spatial Attention Changes Excitability of Human Visual Cortex to Direct Stimulation. Current Biology 17, 134–139 (2007).

92. Haegens, S., Nácher, V., Luna, R., Romo, R. & Jensen, O. α-Oscillations in the monkey sensorimotor network influence discrimination performance by rhythmical inhibition of neuronal spiking. Proc. Natl. Acad. Sci. U. S. A. 108, 19377–19382 (2011).

93. Jensen, O. & Mazaheri, A. Shaping functional architecture by oscillatory alpha activity: Gating by inhibition. Front. Hum. Neurosci. 4, 186 (2010).

94. Hanslmayr, S., Staudigl, T. & Fellner, M.-C. Oscillatory power decreases and long-term memory: The information via desynchronization hypothesis. Front. Hum. Neurosci. 6, 74 (2012).

95. Romei, V., Rihs, T., Brodbeck, V. & Thut, G. Resting electroencephalogram alpha-power over posterior sites indexes baseline visual cortex excitability. Neuroreport 19, 203–208 (2008).

96. Molnar-Szakacs, I. & Uddin, L. Q. Anterior insula as a gatekeeper of executive control. Neurosci. Biobehav. Rev. 139, 104736 (2022).

97. Uddin, L. Q. Salience processing and insular cortical function and dysfunction. Nature Reviews Neuroscience vol. 16 55–61 Preprint at 10.1038/nrn3857 (2015).

98. Vazey, E. M., Moorman, D. E. & Aston-Jones, G. Phasic locus coeruleus activity regulates cortical encoding of salience information. Proc. Natl. Acad. Sci. U. S. A. 115, E9439–E9448 (2018).

99. Nieuwenhuis, S., Aston-Jones, G. & Cohen, J. D. Decision making, the P3, and the locus coeruleus-norepinephrine system. Psychol. Bull. 131, 510–532 (2005).

100. Dahl, M. J., Mather, M. & Werkle-Bergner, M. Noradrenergic modulation of rhythmic neural activity shapes selective attention. Trends Cogn. Sci. 26, 38–52 (2022).

101. Dahl, M. J., Mather, M., Sander, M. C. & Werkle-Bergner, M. Noradrenergic responsiveness supports selective attention across the adult lifespan. Journal of Neuroscience 40, 4372–4390 (2020).

102. Uematsu, A. et al. Modular organization of the brainstem noradrenaline system coordinates opposing learning states. Nat. Neurosci. 20, 1602–1611 (2017).

103. Reid, A. T. et al. Advancing functional connectivity research from association to causation. Nature Neuroscience 2019 22:11 22, 1751–1760 (2019).

104. Beliveau, V. et al. A High-Resolution In Vivo Atlas of the Human Brain’s Serotonin System. The Journal of Neuroscience 37, 120 (2017).

105. Salvi, V. et al. Cingulate cortex stimulation drives distinct pupillary responses in rat via recruitment of noradrenergic neurons in the locus coeruleus. Cerebral Cortex 35, (2025).

106. Molnar-Szakacs, I. & Uddin, L. Q. Anterior insula as a gatekeeper of executive control. Neurosci. Biobehav. Rev. 104736 (2022) doi:10.1016/J.NEUBIOREV.2022.104736.

107. Sara, S. J. & Bouret, S. Orienting and reorienting: The locus coeruleus mediates cognition through arousal. Neuron 76, 130–141 (2012).

108. Gogolla, N. The insular cortex. Current Biology vol. 27 R580–R586 Preprint at 10.1016/j.cub.2017.05.010 (2017).

109. Seeley, W. W. The Salience Network: A Neural System for Perceiving and Responding to Homeostatic Demands. Journal of Neuroscience 39, 9878–9882 (2019).

110. Hermans, E. J. et al. Stress-related noradrenergic activity prompty large-scale neural network reconfiguration. Science (1979). 334, 1151–1153 (2011).

111. Breton-Provencher, V., Drummond, G. T., Feng, J., Li, Y. & Sur, M. Spatiotemporal dynamics of noradrenaline during learned behaviour. Nature 2022 1–7 (2022) doi:10.1038/s41586-022-04782-2.

112. Smegal, L. F. et al. Lower locus coeruleus integrity is associated with diminished practice effects in clinically unimpaired older individuals. Neurobiol. Aging 152, 13–24 (2025).

113. Cleeremans, A., Destrebecqz, A. & Boyer, M. Implicit learning: News from the front. Trends Cogn. Sci. 2, 406–416 (1998).

114. Reber, A. S. Implicit Learning: Background, History, Theory. The Cognitive Unconscious: The First Half Century 3–21 (2022) doi:10.1093/OSO/9780197501573.003.0001.

115. Yu, L. Q., Wilson, R. C. & Nassar, M. R. Adaptive learning is structure learning in time. Neurosci. Biobehav. Rev. 128, 270–281 (2021).

116. Turk-Browne, N. B., Scholl, B. J., Johnson, M. K. & Chun, M. M. Implicit Perceptual Anticipation Triggered by Statistical Learning. Journal of Neuroscience 30, 11177–11187 (2010).

117. Perruchet, P. & Pacton, S. Implicit learning and statistical learning: one phenomenon, two approaches. Trends Cogn. Sci. 10, 233–238 (2006).

118. Udden, J., Folia, V. & Petersson, K. M. The Neuropharmacology of Implicit Learning. Curr. Neuropharmacol. 8, 367–381 (2010).

119. Han, Y. C. et al. Cognitive Neuroscience of Implicit Learning: Implications for Complex Learning and Expertise. The Cognitive Unconscious: The First Half Century 37–61 (2022) doi:10.1093/OSO/9780197501573.003.0003.

120. Howard, D. V. & Howard, J. H. Implicit Learning in Healthy Aging: Evidence from Probabilistic Sequence Learning. The Cognitive Unconscious: The First Half Century 139–154 (2022) doi:10.1093/OSO/9780197501573.003.0007.

121. Jahn, C. I. et al. Dual contributions of noradrenaline to behavioural flexibility and motivation. Psychopharmacology (Berl). 235, 2687–2702 (2018).

122. Dahl, M. J. et al. Rostral locus coeruleus integrity is associated with better memory performance in older adults. Nat. Hum. Behav. 3, 1203–1214 (2019).

123. Lindenberger, U. & Mayr, U. Cognitive aging: Is there a dark side to environmental support? Trends Cogn. Sci. 18, 7–15 (2014).

124. Craik, F. I. M. On the transfer of information from temporary to permanent memory. Philosophical Transactions of the Royal Society B: Biological Sciences 302, 341–359 (1983).

125. Briand, L. A., Gritton, H., Howe, W. M., Young, D. A. & Sarter, M. Modulators in concert for cognition: Modulator interactions in the prefrontal cortex. Prog. Neurobiol. 83, 69–91 (2007).

126. Bangasser, D. A., Wiersielis, K. R. & Khantsis, S. Sex differences in the locus coeruleus-norepinephrine system and its regulation by stress. Brain Res. 1641, 177–188 (2016).

127. Merz, C. J., Kinner, V. L. & Wolf, O. T. Let’s talk about sex… differences in human fear conditioning. Curr. Opin. Behav. Sci. 23, 7–12 (2018).

128. Gruene, T. M., Flick, K., Stefano, A., Shea, S. D. & Shansky, R. M. Sexually divergent expression of active and passive conditioned fear responses in rats. Elife 4, (2015).

129. Voulo, M. E. & Parsons, R. G. Response-specific sex difference in the retention of fear extinction. Learn. Mem. 24, 245–251 (2017).

130. Mulvey, B. et al. Molecular and Functional Sex Differences of Noradrenergic Neurons in the Mouse Locus Coeruleus. Cell Rep. 23, 2225–2235 (2018).

131. Duszkiewicz, A. J., McNamara, C. G., Takeuchi, T. & Genzel, L. Novelty and dopaminergic modulation of memory persistence: A tale of two systems. Trends Neurosci. 42, 102–114 (2019).

132. Kret, M. E. & Sjak-Shie, E. E. Preprocessing pupil size data: Guidelines and code. Behav. Res. Methods 51, 1336–1342 (2019).

133. Dahl, M. J. et al. Rostral locus coeruleus integrity is associated with better memory performance in older adults. Nat. Hum. Behav. 3, 1203–1214 (2019).

134. Fernandes, P., Regala, J., Correia, F. & Gonçalves-Ferreira, A. J. The human locus coeruleus 3-D stereotactic anatomy. Surgical and Radiologic Anatomy 34, 879–885 (2012).

135. Prokopiou, P. C. et al. Lower novelty-related locus coeruleus function is associated with Aβ-related cognitive decline in clinically healthy individuals. Nat. Commun. 13, (2022).

136. Lee, T.-H. et al. Arousal increases neural gain via the locus coeruleus–noradrenaline system in younger adults but not in older adults. Nat. Hum. Behav. 2, 356–366 (2018).

137. Halchenko, Y. O. et al. HeuDiConv — flexible DICOM conversion into structured directory layouts. J. Open Source Softw. 9, 5839 (2024).

138. Esteban, O. et al. Crowdsourced MRI quality metrics and expert quality annotations for training of humans and machines. Sci. Data 6, 30 (2019).

139. Esteban, O. et al. MRIQC: Advancing the automatic prediction of image quality in MRI from unseen sites. PLoS One 12, e0184661 (2017).

140. Esteban, O. et al. fMRIPrep: a robust preprocessing pipeline for functional MRI. Nat. Methods 16, 111–116 (2019).

141. Worsley, K. J. & Friston, K. J. Analysis of fMRI Time-Series Revisited—Again. Neuroimage 2, 173–181 (1995).

142. Satterthwaite, T. D. et al. An improved framework for confound regression and filtering for control of motion artifact in the preprocessing of resting-state functional connectivity data. Neuroimage 64, 240–256 (2013).

143. Friston, K. J., Williams, S., Howard, R., Frackowiak, R. S. J. & Turner, R. Movement-Related effects in fMRI time-series. Magn. Reson. Med. 35, 346–355 (1996).

144. Delorme, A. & Makeig, S. EEGLAB: An open source toolbox for analysis of single-trial EEG dynamics including independent component analysis. J. Neurosci. Methods 134, 9–21 (2004).

145. Dimigen, O., Sommer, W., Hohlfeld, A., Jacobs, A. M. & Kliegl, R. Coregistration of eye movements and EEG in natural reading: Analyses and review. J. Exp. Psychol. Gen. 140, 552–572 (2011).

146. Oostenveld, R., Fries, P., Maris, E. & Schoffelen, J. M. FieldTrip: Open source software for advanced analysis of MEG, EEG, and invasive electrophysiological data. Comput. Intell. Neurosci. 2011, 1–9 (2011).

147. Nolan, H., Whelan, R. & Reilly, R. B. FASTER: Fully Automated Statistical Thresholding for EEG artifact Rejection. J. Neurosci. Methods 192, 152–162 (2010).

148. Maris, E. & Oostenveld, R. Nonparametric statistical testing of EEG- and MEG-data. J. Neurosci. Methods 164, 177–190 (2007).

149. Friston, K. J., Ashburner, J., Kiebel, S., Nichols, T. & Penny, W. Statistical Parametric Mapping: The Analysis of Functional Brain Images. Statistical Parametric Mapping: The Analysis of Functional Brain Images (Academic Press, Cambridge, Massachusetts, 2007). doi:10.1016/B978-0-12-372560-8.X5000-1.

150. Friston, K. J. et al. Event-Related fMRI: Characterizing Differential Responses. Neuroimage 7, 30–40 (1998).

151. Smith, S. M. et al. Network modelling methods for FMRI. Neuroimage 54, 875–891 (2011).

152. Avants, B. B., Tustison, N. & Song, G. Advanced Normalization Tools: V1.0. Insight Journal 2, (2009).

153. Tustison, N. J. et al. The ANTsX ecosystem for quantitative biological and medical imaging. Scientific Reports 2021 11:1 11, 1–13 (2021).

154. Murphy, P. R., O’Connell, R. G., O’Sullivan, M., Robertson, I. H. & Balsters, J. H. Pupil diameter covaries with BOLD activity in human locus coeruleus. Hum. Brain Mapp. 35, 4140–4154 (2014).

155. Theiler, J., Eubank, S., Longtin, A., Galdrikian, B. & Doyne Farmer, J. Testing for nonlinearity in time series: the method of surrogate data. Physica D 58, 77–94 (1992).

156. Krishnan, A., Williams, L. J., McIntosh, A. R. & Abdi, H. Partial Least Squares (PLS) methods for neuroimaging: A tutorial and review. Neuroimage 56, 455–475 (2011).

157. McIntosh, A. R. & Lobaugh, N. J. Partial least squares analysis of neuroimaging data: Applications and advances. in NeuroImage vol. 23 (Neuroimage, 2004).

158. Brett, M., Anton, J.-L., Valabregue, R. & Poline, J.-B. Region of interest analysis using an SPM toolbox. In Human Brain Mapping conference (2002).

