## Supplementary methods and results for "Locus coeruleus-related insula activation is associated with implicit learning"

*** Corresponding author**:

**ORCIDs:**

0000-0003-3525-0539
0009-0004-8578-1092
0000-0003-4331-6112
0000-0002-6399-9996

### 1. Supplementary methods

Results included in this manuscript come from preprocessing performed using *fMRIPrep* 20.0.1 (Esteban, Markiewicz, et al. (2018); Esteban, Blair, et al. (2018); RRID:SCR_016216), which is based on *Nipype* 1.4.2 (Gorgolewski et al. (2011); Gorgolewski et al. (2018); RRID:SCR_002502).

#### Anatomical data preprocessing

The T1-weighted (T1w) image was corrected for intensity non-uniformity (INU) with N4BiasFieldCorrection (Tustison et al. 2010), distributed with ANTs 2.2.0 (Avants et al. 2008, RRID:SCR_004757), and used as T1w-reference throughout the workflow. The T1w-reference was then skull-stripped with a *Nipype* implementation of the antsBrainExtraction.sh workflow (from ANTs), using OASIS30ANTs as target template. Brain tissue segmentation of cerebrospinal fluid (CSF), white-matter (WM) and gray-matter (GM) was performed on the brain-extracted T1w using fast (FSL 5.0.9, RRID:SCR_002823, Zhang, Brady, and Smith 2001). Brain surfaces were reconstructed using recon-all (FreeSurfer 6.0.1, RRID:SCR_001847, Dale, Fischl, and Sereno 1999), and the brain mask estimated previously was refined with a custom variation of the method to reconcile ANTs-derived and FreeSurfer-derived segmentations of the cortical gray-matter of Mindboggle (RRID:SCR_002438, Klein et al. 2017). Volume-based spatial normalization to two standard spaces (MNI152NLin2009cAsym, MNI152NLin6Asym) was performed through nonlinear registration with antsRegistration (ANTs 2.2.0), using brain-extracted versions of both T1w reference and the T1w template. The following templates were selected for spatial normalization: *ICBM 152 Nonlinear Asymmetrical template version 2009c* [Fonov et al. (2009), RRID:SCR_008796; TemplateFlow ID: MNI152NLin2009cAsym], *FSL's MNI ICBM 152 non-linear 6th Generation Asymmetric Average Brain Stereotaxic Registration Model* [Evans et al. (2012), RRID:SCR_002823; TemplateFlow ID: MNI152NLin6Asym],

#### Functional data preprocessing

For each of the 1 BOLD runs found per subject (across all tasks and sessions), the following preprocessing was performed. First, a reference volume and its skull-stripped version were generated using a custom methodology of *fMRIPrep*. A deformation field to correct for susceptibility distortions was estimated based on *fMRIPrep*'s *fieldmap-less* approach. The deformation field is that resulting from co-registering the BOLD reference to the same-subject T1w-reference with its intensity inverted (Wang et al. 2017; Huntenburg 2014). Registration is performed with antsRegistration (ANTs 2.2.0), and the process regularized by constraining deformation to be nonzero only along the phase-encoding direction, and modulated with an average fieldmap template (Treiber et al. 2016). Based on the estimated susceptibility distortion, a corrected EPI (echo-planar imaging) reference was calculated for a more accurate co-registration with the anatomical reference. The BOLD reference was then co-registered to the T1w reference using bbregister (FreeSurfer) which implements boundary-based registration (Greve and Fischl 2009). Co-registration was configured with six degrees of freedom. Head-motion parameters with respect to the BOLD reference (transformation matrices, and six corresponding rotation and translation parameters) are estimated before any spatiotemporal filtering using mcflirt (FSL 5.0.9, Jenkinson et al. 2002). BOLD runs were slice-time corrected using 3dTshift from AFNI 20160207 (Cox and Hyde 1997, RRID:SCR_005927). The BOLD time-series were resampled onto the following surfaces (FreeSurfer reconstruction nomenclature): *fsnative*, *fsaverage*. The BOLD time-series (including slice-timing correction when applied) were resampled onto their original, native space by applying a single, composite transform to correct for head-motion and susceptibility distortions. These resampled BOLD time-series will be referred to as *preprocessed BOLD in* *original space*, or just *preprocessed BOLD*. The BOLD time-series were resampled into standard space, generating a *preprocessed BOLD run in MNI152NLin2009cAsym space*. First, a reference volume and its skull-stripped version were generated using a custom methodology of *fMRIPrep*. Automatic removal of motion artifacts using independent component analysis (ICA-AROMA, Pruim et al. 2015) was performed on the *preprocessed BOLD on MNI space* time-series after removal of non-steady state volumes and spatial smoothing with an isotropic, Gaussian kernel of 6mm FWHM (full-width half-maximum). Corresponding "non-aggresively" denoised runs were produced after such smoothing. Additionally, the "aggressive" noise-regressors were collected and placed in the corresponding confounds file. Several confounding time-series were calculated based on the *preprocessed BOLD*: framewise displacement (FD), DVARS and three region-wise global signals. FD and DVARS are calculated for each functional run, both using their implementations in *Nipype* (following the definitions by Power et al. 2014). The three global signals are extracted within the CSF, the WM, and the whole-brain masks. Additionally, a set of physiological regressors were extracted to allow for component-based noise correction (*CompCor*, Behzadi et al. 2007). Principal components are estimated after high-pass filtering the *preprocessed BOLD* time-series (using a discrete cosine filter with 128s cut-off) for the two *CompCor* variants: temporal (tCompCor) and anatomical (aCompCor). tCompCor components are then calculated from the top 5% variable voxels within a mask covering the subcortical regions. This subcortical mask is obtained by heavily eroding the brain mask, which ensures it does not include cortical GM regions. For aCompCor, components are calculated within the intersection of the aforementioned mask and the union of CSF and WM masks calculated in T1w space, after their projection to the native space of each functional run (using the inverse BOLD-to-T1w transformation). Components are also calculated separately within the WM and CSF masks. For each CompCor decomposition, the *k* components with the largest singular values are retained, such that the retained components' time series are sufficient to explain 50 percent of variance across the nuisance mask (CSF, WM, combined, or temporal). The remaining components are dropped from consideration. The head-motion estimates calculated in the correction step were also placed within the corresponding confounds file. The confound time series derived from head motion estimates and global signals were expanded with the inclusion of temporal derivatives and quadratic terms for each (Satterthwaite et al. 2013). Frames that exceeded a threshold of 0.5 mm FD or 1.5 standardised DVARS were annotated as motion outliers. All resamplings can be performed with *a single interpolation step* by composing all the pertinent transformations (i.e. head-motion transform matrices, susceptibility distortion correction when available, and co-registrations to anatomical and output spaces). Gridded (volumetric) resamplings were performed using antsApplyTransforms (ANTs), configured with Lanczos interpolation to minimize the smoothing effects of other kernels (Lanczos 1964). Non-gridded (surface) resamplings were performed using mri_vol2surf (FreeSurfer).

Many internal operations of *fMRIPrep* use *Nilearn* 0.6.2 (Abraham et al. 2014, RRID:SCR_001362), mostly within the functional processing workflow. For more details of the pipeline, see the section corresponding to workflows in *fMRIPrep*'s documentation.

**Copyright waiver**

The above boilerplate text was automatically generated by fMRIPrep with the express intention that users should copy and paste this text into their manuscripts *unchanged*. It is released under the CC0 license.

#### 2. Supplementary results

##### Pupil–BOLD associations with variable temporal delays

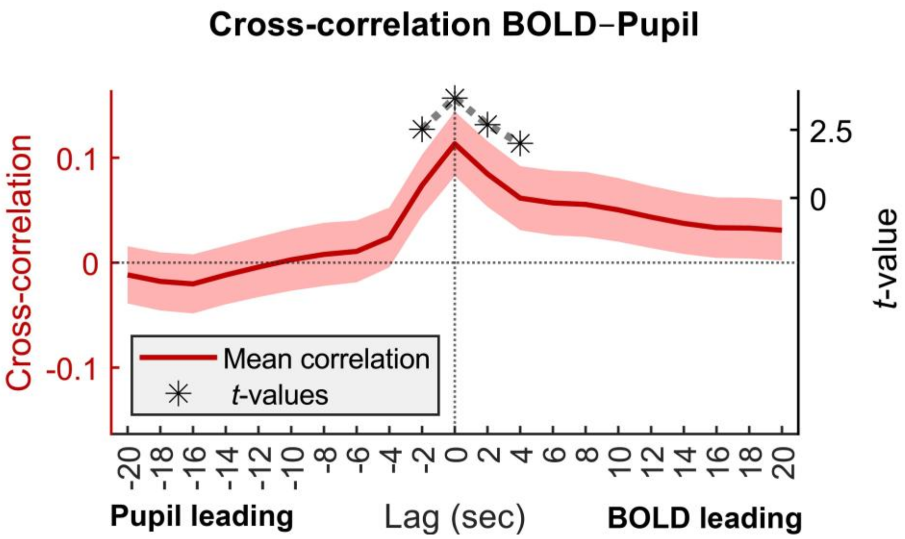

***Figure S1*. Cross-correlation of BOLD activity and pupil size.** BOLD activity of regions showing a reliable association with pupil size (cf. main text Figure 4) was averaged and correlated with moment-to-moment changes in pupil size for each participant. A group-level cluster-based random permutation test evaluated the association across participants while controlling for multiple comparisons. *t*-values for lags (–2)–(+4): 2.520, 3.663, 2.689, 2.004. For all other lags, *t*-values fall below 1.96.

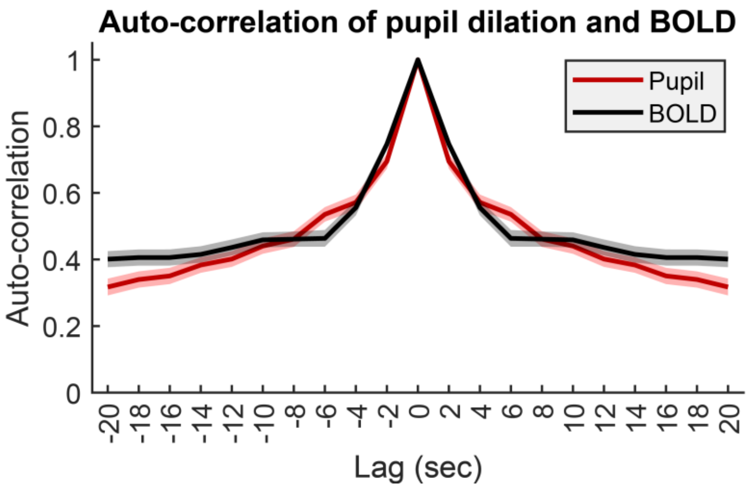

***Figure S2*. Auto-correlation of BOLD activity and pupil size.** BOLD activity of regions showing a reliable association with pupil size (cf. main text Figure 4) was averaged. Then the auto-correlation of these regions and of moment-to-moment changes in pupil size was compared. The two signals exhibit comparable temporal persistence — their fluctuations evolve on similar timescales.

**Brainstem alignment quality control**

To guarantee accurate analyses of small subcortical structures, brainstem alignment was quantified using a landmark at the pons–fourth ventricle boundary located at the rostrocaudal level of the locus coeruleus (Figure S3a). For each participant, intensity profiles surrounding this landmark were extracted from the warped mean EPI image and compared with the MNI template (15 data points; standardized intensities with flipped sign to match contrast across modalities). Warped EPIs closely matched the MNI template's intensity pattern (median correlation coefficients across participants $r = 0.817$; Figure S3b). Artificially shifting participants' warped EPI images by 1–2 voxels in XYZ direction markedly reduced correspondence (median $r = 0.272$, $p < 0.001$), indicating that this metric was sensitive to identify misregistration.

To test robustness, pupil–BOLD analyses were repeated after excluding participants with low brainstem alignment quality (correlation < 0.5; cf. ; Figure S3c) or insufficient valid pupil data (<50% samples; see main text *Methods* section). The resulting group maps were highly similar to the full-sample results (voxel-wise *t*-map correlation $r = 0.932$), with a tendency of the more stringent participant exclusion to *enhance* both positive and negative pupil–BOLD associations (Figure S3d). Together, this indicates that the reported findings are unlikely driven by poor registration or pupillometry quality.

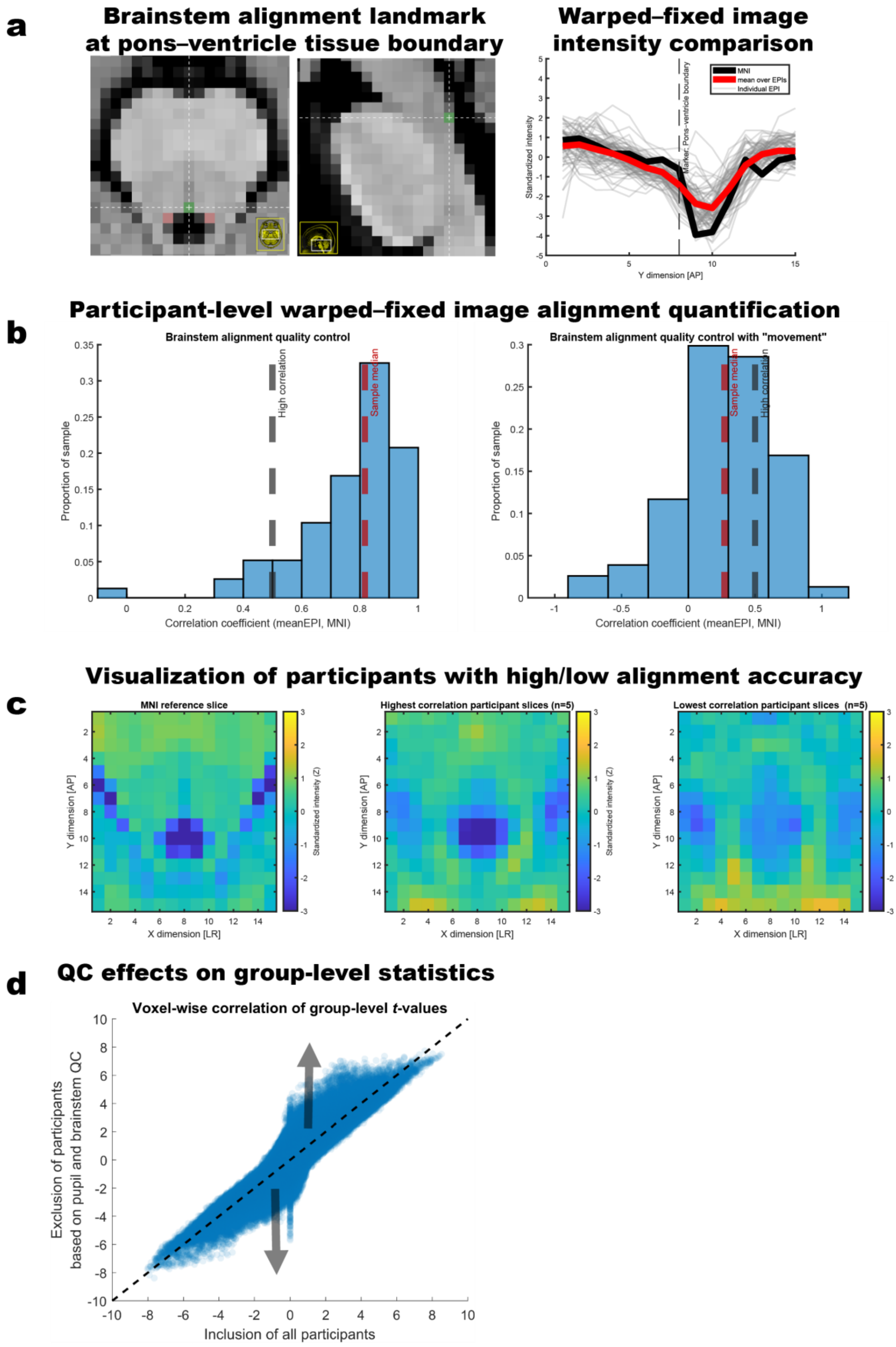

***Figure S3*. Landmark-based quantification of brainstem alignment accuracy (a) is sensitive to movement (b–c) and increases group-level pupil–BOLD associations (d).** QC, quality control. EPI, Echo,planar imaging. MNI, Montreal Neurological Institute (MNI) standard space.

**Pupil–locus coeruleus BOLD associations**

After verifying accurate alignment, we additionally performed anatomy-informed locus coeruleus region-of-interest analyses. For this, the locus coeruleus mask was slightly dilated to accommodate the lower fMRI resolution (2 mm FWHM smoothing and thresholding at 0.001; excluding fourth-ventricle voxels) in keeping with earlier research. These analyses revealed significant pupil–BOLD associations spatially overlapping with or in direct proximity to the locus coeruleus for all tested comparisons.

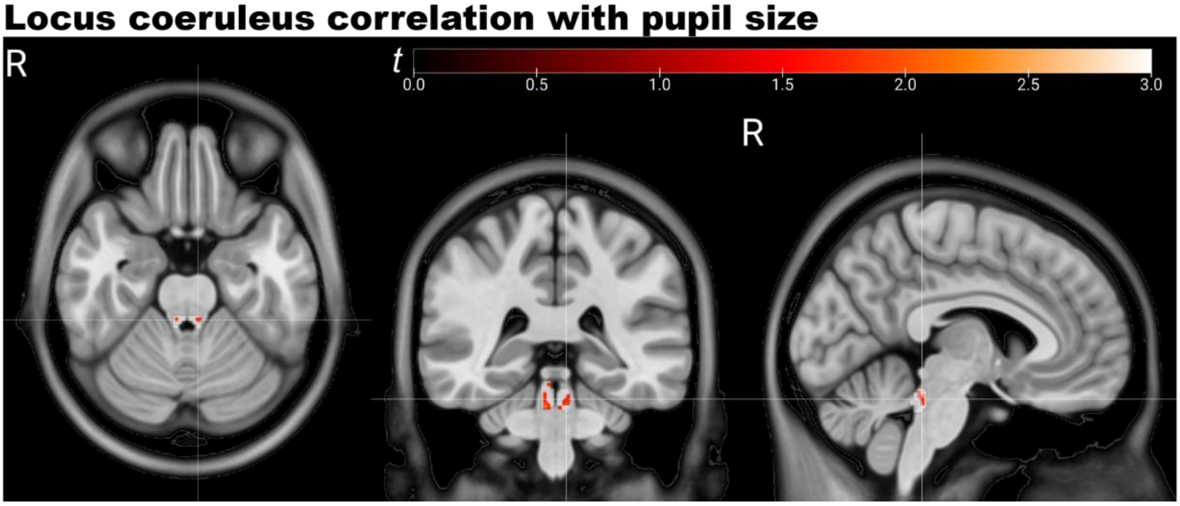

***Figure S4*****. Moment-to-moment changes in pupil size correlate with locus coeruleus BOLD signals.** Example visualization of locus coeruleus–BOLD association for subsample with high quality brainstem alignment and pupil data as well as minimal smoothing (2 mm FWHM). The white background area indicates the applied locus coeruleus region of interest. The overlaid group-level result map (*t*-values) is thresholded at p < 0.05.

**Table S1:** Pupil–locus coeruleus–associations across subsamples and fMRI smoothing kernels

| **Sample description** | **Smoothing (FWHM)** | ***df*** | ***t*** | ***p*** | **MNI X Y Z (mm)** |
|---|---|---|---|---|---|
| Full sample,default smoothing | 8 | 70 | 2.97 | 0.002 | 6 –34 –16 |
| Good alignment + pupil subsample, default smoothing | 8 | 57 | 3.77 | <0.001 | 2 –34 –16 |
| Full sample, narrow smoothing | 2 | 70 | 3.06 | 0.001 | –6 –36 –16 |
| Good alignment + pupil subsample, narrow smoothing | 2 | 57 | 3.36 | 0.001 | 2 –36 –24 |

**Pupil–locus coeruleus associations with control for dopaminergic neuromodulation**

We additionally investigated whether controlling for BOLD signals in the substantia nigra and ventral tegmental area would abolish the pupil–locus coeruleus BOLD association, using a voxel-wise multiple regression framework (masks derived from: https://www.nitrc.org/projects/brainstemnavig). In short, for each participant, for each voxel falling into our locus coeruleus mask we estimated the association of moment-to-moment changes in pupil and BOLD signals, while controlling for nuisance variables (cf. main text *Methods* section) and also BOLD signals averaged over substantia nigra and ventral tegmental area voxels. Next, on a group level we compared participant-level beta parameters for the pupil–locus coeruleus BOLD association estimated from analyses with and without substantia nigra–ventral tegmental area covariate. While there was a slight numerical attenuation of the association when including the covariate, for no voxel this reached significance (dependent samples *t*-test, all absolute *t*-values < 1) when using minimally smoothed fMRI data. Together, this indicates that the association between pupil and locus coeruleus BOLD signals is not explained by the co-activation of dopaminergic structures when narrow smoothing kernels are used. However, for larger smoothing kernels, covariate inclusion attenuated the pupil–BOLD association, indicating a dependency of signals. In line with previous work, we assume that multiple neuromodulatory systems contribute to pupil changes.

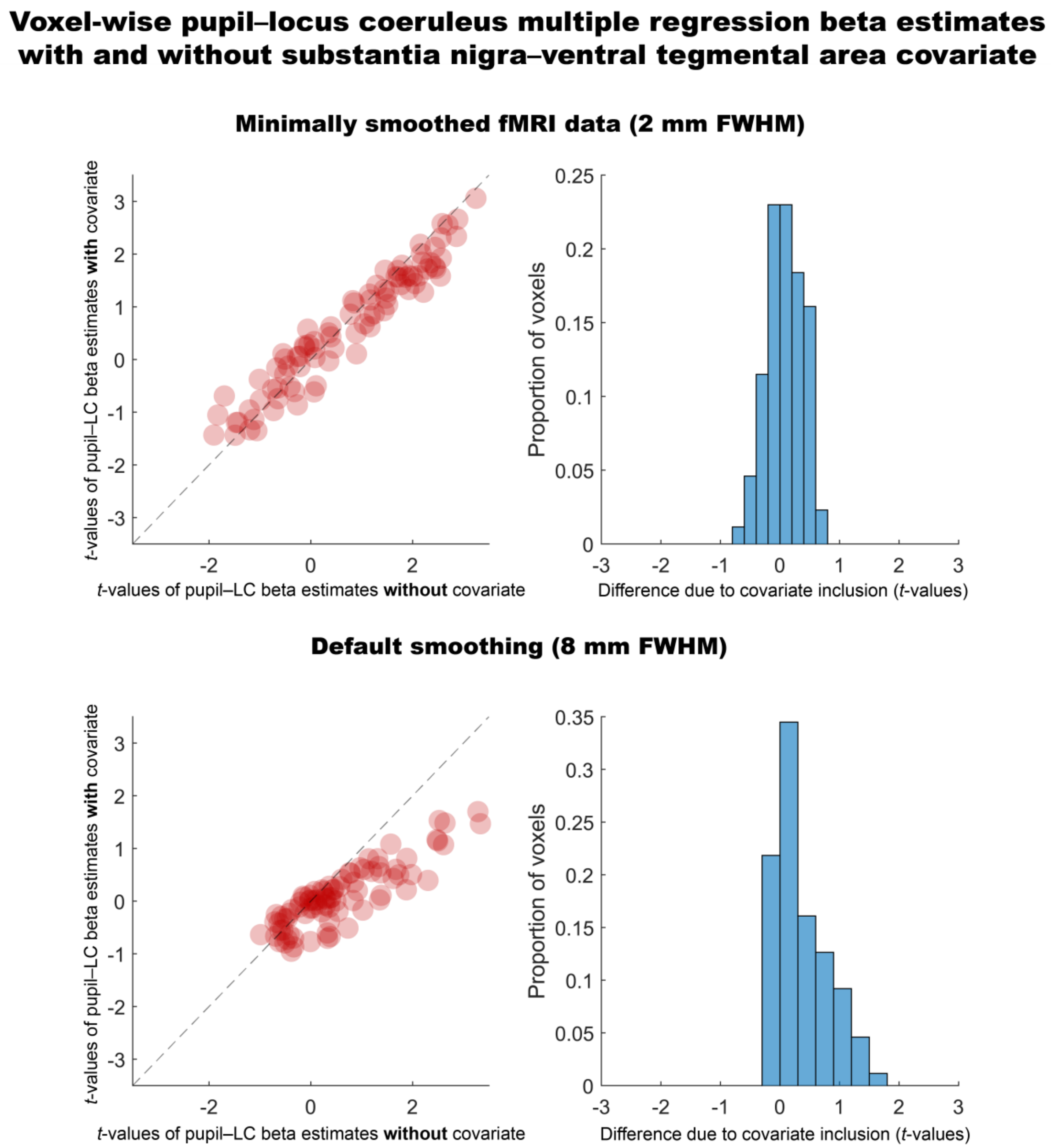

***Figure S5***. **Moment-to-moment changes in pupil size correlate with locus coeruleus BOLD signals even after inclusion of dopaminergic covariates when data are minimally smoothed.**

#### Pupil- and locus coeruleus- associations with default-mode network BOLD signals

**Table S2: P**upil–default-mode network associations across subsamples and smoothing kernels

| Sample description | Smoothing (FWHM) | *df* | *t* | *p* |
|---|---|---|---|---|
| Full sample, default smoothing | 8 | 70 | -2.165 | 0.034 |
| Good alignment + pupil subsample, default smoothing | 8 | 57 | -2.711 | 0.009 |
| Full sample, narrow smoothing | 2 | 70 | -1.541 | 0.128 |
| Good alignment + pupil subsample, narrow smoothing | 2 | 57 | -2.282 | 0.026 |

*Note*: Cortical default-mode network BOLD signals were extracted and averaged based on the Yeo et al. (2011) atlas available through Lead-DBS (https://www.lead-dbs.org/).

**Table S3: Locus coeruleus**–default-mode network associations across subsamples and smoothing kernels

| Sample description | Smoothing (FWHM) | *df* | *t* | *p* |
|---|---|---|---|---|
| Full sample, default smoothing | 8 | 70 | -4.838 | <0.001 |
| Good alignment + pupil subsample, default smoothing | 8 | 57 | -5.118 | <0.001 |
| Full sample, narrow smoothing | 2_8 | 70 | -3.211 | 0.002 |
| Good alignment + pupil subsample, narrow smoothing | 2_8 | 57 | -2.925 | 0.005 |

*Note*: Cortical default-mode network BOLD signals were extracted and averaged based on the Yeo et al. (2011) atlas available through Lead-DBS (https://www.lead-dbs.org/). Smoothing 2_8 indicates a narrow smoothing for the locus coeruleus seed region and a default (8 mm) smoothing for target voxels.

213 **PET transporter distribution overlaps with pupil-linked BOLD activation patterns**

214 **Table S4:** Variance explained in pupil–BOLD maps by PET-derived neuromodulatory transporter maps

| Model | Ordinary R-squared (unadjusted) | R-squared adjusted for the number of coefficients |
|---|---|---|
| Base (DAT, 5HTT) | 0.085 | 0.085 |
| Full (NAT, DAT, 5HTT)) | 0.099 | 0.099 |

215 *Note*: NAT, noradrenergic transporter; DAT, dopaminergic transporter; 5-HTT, serotonergic transporter

216

#### Dynamic Causal Modeling-based locus coeruleus–anterior insula connectivity analysis

For all participants, fully connected DCM models showed substantially improved model evidence compared with a null model lacking inter-regional connections and modulatory effects (Figure S6).

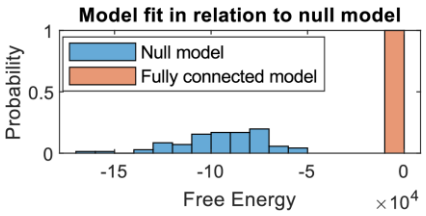

**Figure S6**. **DCM model fit of fully connected and null models** (shown for 2 mm FWHM smoothed data).

Group-level Parametric Empirical Bayes (PEB) analyses revealed reliable intrinsic connectivity between the locus coeruleus and anterior insula. The winning reduced model consistently retained an excitatory locus coeruleus→anterior insula connection (for narrow 2 mm and default 8 mm fMRI smoothing kernels). Evidence for the reciprocal anterior insula→locus coeruleus connection was present only for the narrow smoothing kernel, suggesting that this better preserves signals of the small brainstem nucleus. Comparing the two connectivity parameters (Bayesian contrast within the PEB framework; 2 mm smoothing data) provided little evidence that the Insula→LC connection differed from the LC→Insula connection (posterior probability = 0.035). Finally, the oddball condition modulated the locus coeruleus self-connection (2 mm smoothing), consistent with a task-related *increase* in locus coeruleus excitability (Figure S7; Table S5).

Together, these results indicate directed functional coupling between anterior insula and locus coeruleus, with a reciprocal influence between the two regions and a task-related modulation of noradrenergic dynamics. The DCM findings converge with and extend the lag-based connectivity results reported in the main text, supporting bidirectional locus coeruleus–insula interactions during salient oddball processing.

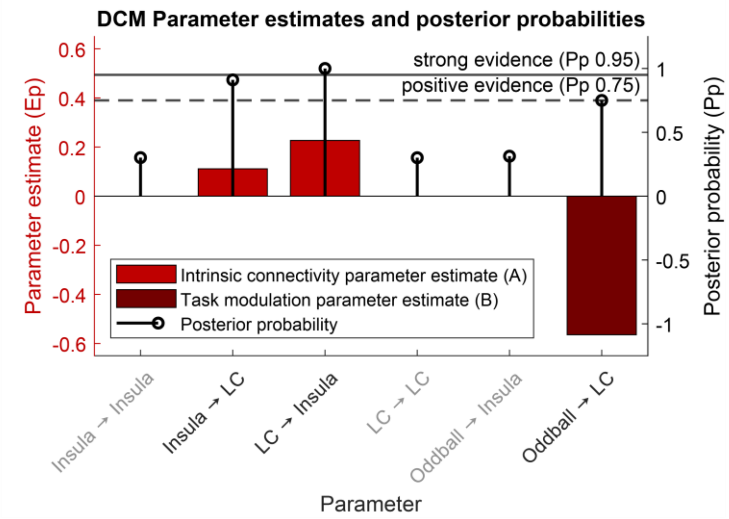

**Figure S7**. **Parameter estimates and posterior probabilities suggesting reliable intrinsic locus coeruleus–anterior insula connectivity and task-related modulation of locus coeruleus excitability.** Results of the DCM estimation (winning Bayesian Model Reduction-pruned Parametric Empirical Bayes model) with 2 mm FWHM smoothed fMRI data are shown. Note that positive inter-regional intrinsic connectivity parameter estimates indicate excitatory influences (A matrix; e.g., LC→Insula), while negative task modulation parameter estimates (B matrix; e.g., Oddball→LC) indicate task-related increases in regional excitability. Greyed out font indicates posterior probability estimates < 0.75 (below positive evidence).

**Table S5: Locus coeruleus–anterior insula connectivity for winning DCM models**

| Smoothing (mm) | Variable | Insula→ Insula | Insula→ LC | LC→ Insula | LC→ LC | Oddball→ Insula | Oddball→ LC |
|---|---|---|---|---|---|---|---|
| 8 | Parameter estimate (Ep) | 0 | -0.043 | **0.186** | 0 | 0 | 0 |
| 8 | Posterior probability (Pp) | 0.307 | 0.561 | **1** | 0.335 | 0.405 | 0.41 |
| 2 | Parameter estimate (Ep) | 0 | **0.111** | **0.227** | 0 | 0 | **-0.565** |
| 2 | Posterior probability (Pp) | 0.302 | **0.911** | **1** | 0.301 | 0.313 | **0.75** |

*Note*: Greyed out path show posterior probability estimates < 0.75 (below positive evidence). Note that positive inter-regional intrinsic connectivity parameter estimates indicate excitatory influences (A matrix; e.g., LC→Insula), while negative task modulation parameter estimates (B matrix; e.g., Oddball→LC) indicate task-related increases in regional excitability. LC, locus coeruleus. Re-estimating participant-level DCM models with mean centered inputs reproduced the positive Insula→LC and LC→Insula connectivity (posterior probabilities > 0.95), but evidence for the Oddball→LC modulation decreased (posterior probability = 0.37; 2 mm smoothing data). For a discussion of the implications of mean-centering for parameter interpretation, see Zeidman et al. (2019).

As sensitivity analysis, we repeated the cross-correlation (main text Figure 7e) and DCM (supplementary information Figure S7) analyses using a study-independent anterior-insula region of interest definition (downloaded from: https://soundray.org/hammers-n30r95/ ; archive: Hammers-newInsula_regions-probmaps-gm+full.tar, file: probmap-gm-insula_anterior_pole_L.nii.gz, probabilistic ROI thresholded at 75%).

Using this alternative approach, the cross-correlation analysis again showed the temporal precedence of anterior insula relative to locus coeruleus BOLD activity (contrast of lags [(–4)–(–2)] vs [(+2)–(+4)]), $t = 2.06$; $p = 0.043$.

The group-level DCM analysis of 2 mm smoothed data again identified a winning model that maintained the directed anterior insula↔locus coeruleus connections: insula→locus coeruleus, parameter estimate (Ep) 0.086, posterior probability (Pp) 0.777; locus coeruleus→insula, parameter estimate (Ep) 0.11, posterior probability (Pp) 0.982.

Together, this sensitivity analysis indicates that our initial cross-correlation and DCM findings (main text Figure 7e, supplementary information Figure S7) are unlikely driven by how we defined the anterior-insula region of interest.

**Response time dynamics over the course of the conditioned oddball task**

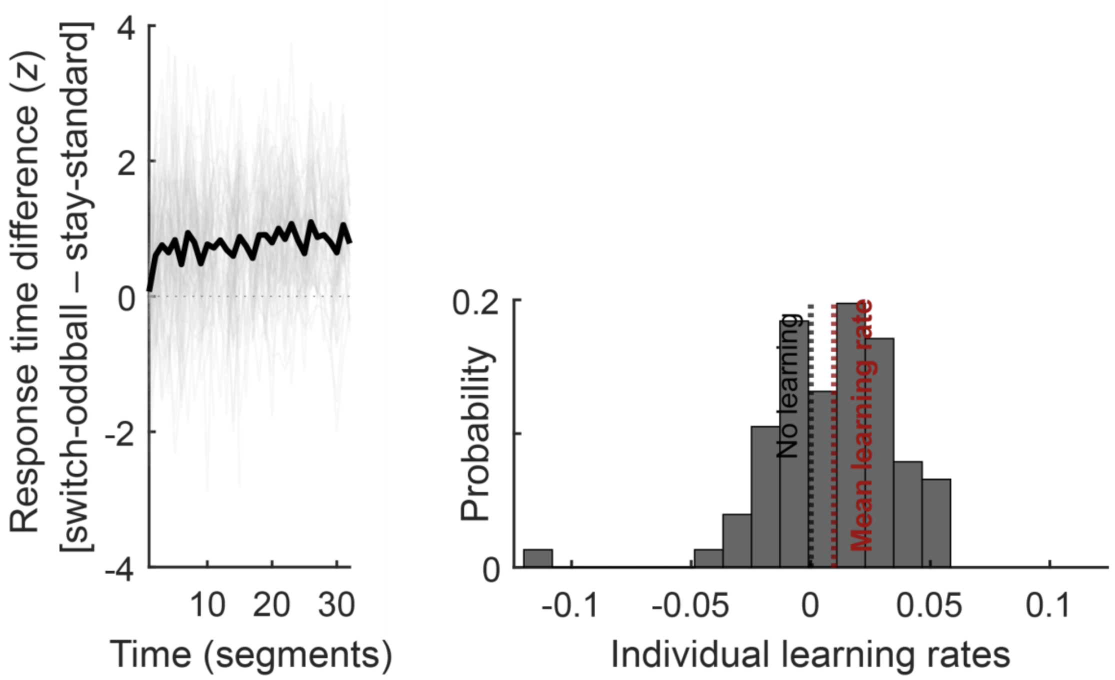

**Figure S8. Participant-level implicit learning curves (left) and slope parameters (right).** Participant focus their behavior on more likely trial categories at the expense of less likely categories. This bias emerges across the course of the experiments (left) indicating implicit learning of the task structure. The mean response time difference is shown in black, while individual participants are shown in light grey. The majority of participants shows a positive linear slope parameter (right; see red mean learning rate line), indicating larger response time differences over the course of the experiments. Note that each time segment corresponds to 10 trials, including both repetitions of the conditioned oddball experiment.

**Table S6:** Response times by trial category over the course of the conditioned oddball task

| **Effect** | **Estimate** | **SE** | ***t*** | ***df*** | ***p*** | **CI_Lower** | **CI_Upper** | **Interpretation** |
|---|---|---|---|---|---|---|---|---|
| Intercept | -0.189 | 0.014 | -13.338 | 2803 | <.001 | -0.217 | -0.161 | Baseline response times (standard condition, mean over trials) |
| Oddball cond. | 0.631 | .042 | 4.892 | 2803 | <.001 | 0.548 | 0.714 | Difference in response times between standard/oddball conditions |
| Trial # | -0.001 | <0.001 | -9.076 | 2803 | <.001 | -0.001 | -0.001 | Overall change in response times across trials (learning effect) |
| Trial# :Oddball cond. | 0.002 | <0.001 | 11.035 | 2803 | <.001 | 0.001 | 0.002 | Difference in learning rate across trials between standard/oddball conditions |

*Note*: Response times were analyzed using a linear mixed-effects model with trial number (mean-centered) and trial condition (standard/oddball) as categorical within-participant predictors. The model included by-participant random intercepts and random slopes for both predictors.

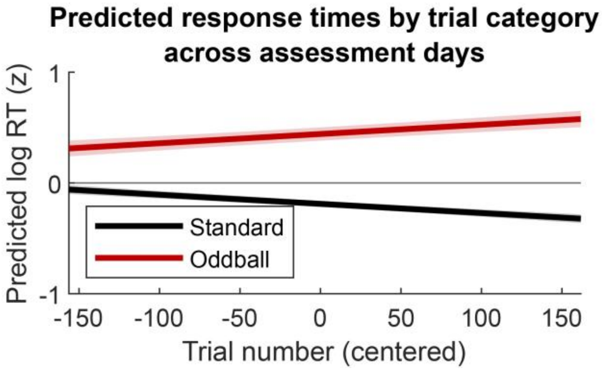

**Figure S9**. **Predicted response times by trial category over assessment days** (based on fixed effects only; cf. Table S6 for model output).

An additional **two-way analysis of variance (ANOVA)** was conducted to examine the participant-level **effects of Response (Stay/Switch) and Trial condition (Standard/Oddball)** on response times (log RT [*z*]).

There was a significant main effect of Response, $F(1, 280) = 123.07$, $p < .001$, indicating that the response times differed when participants repeated the same response vs. switched to another response button (marginal means ± standard error: 0.003 ± 0.025, 0.387 ± 0.025). There was also a significant main effect of Trial condition, $F(1, 280) = 356.26$, $p < .001$, indicating that standard and oddball trials differed in their response times(marginal means ± standard error: -0.132 ± 0.025, 0.522 ± 0.025). Finally, the interaction between the factors Response and Trial condition was significant, $F(1, 280) = 42.15$, $p < .001$, suggesting that the effect of the factor Response on response times depended on the level of Trial condition.

#### Response times and physiological responses by trial category

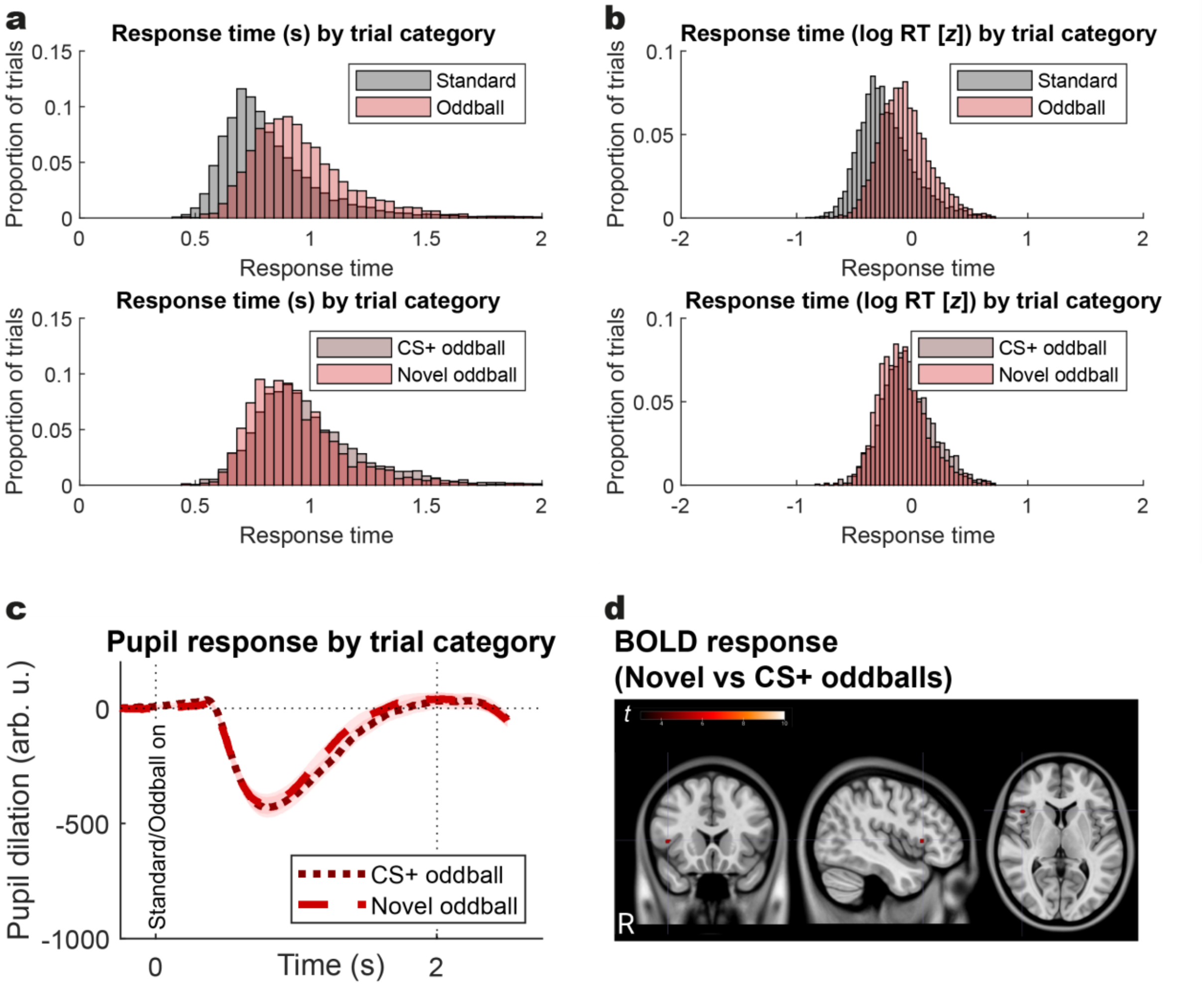

**Figure S10. Response times ((a) raw, (b) log transformed and *z*-scored), (c) pupil and (d) BOLD responses for novel oddball stimuli as well as previously fear conditioned oddball stimuli**. **(a-b)** Novel compared to previously fear-conditioned oddball stimuli are associated with slightly quicker response times (also see Table S7). **(c)** Task-related pupil responses to novel and previously fear conditioned oddball stimuli are comparable (group-level cluster-based random permutation test: $p_{cluster\text{-}corr} \geq 0.08$). **(d)** Task-related BOLD responses to novel and previously fear conditioned oddball stimuli show spatially constrained differences (see crosshair; CS+>novel: $z = 5$, $p_{FWE\text{-}corr} \leq 0.001$; MNI: 43.5, 19.5, 5.5 [right frontal operculum]. For visualization, the group contrast maps are thresholded at $p_{FWE\text{-}corr} \leq 0.05$).

**Table S7:** Linear mixed effects model contrasting CS+ and novel oddball subtypes

| Outcome | Estimate | SE | *t* | *df* | *p* | CI_Lower | CI_Upper |
|---|---|---|---|---|---|---|---|
| Response time (log RT [*z*]) | -0.193 | 0.05 | -3.853 | 3385 | <.001 | -0.292 | -0.095 |

*Note*: Trial-level linear mixed effects models probing if CS+ oddballs differed systematically from novel oddballs in terms of response times (log RT [z]), using the following model formula: RT ~ oddball-category + age-group + (1+ oddball-category | ID)

#### Brain–behavior analyses split by neural indicator and age group comparisons

**Table S8:** Associations of individual neural indicators with implicit learning (latent behavioral scores)

| Variable | nYA, nOA | *df* | *rho* | *p* |
|---|---|---|---|---|
| Locus coeruleus integrity | 36, 32 | 68 | 0.224 | 0.067 |
| Pupil dilation | 36, 32 | 66 | 0.207 | 0.090 |
| Insula BOLD | 36, 32 | 66 | 0.356 | 0.003 |

*Note*: The association of each neural indicator with the latent behavioral implicit learning scores was evaluated. See Methods section (partial least squares analyses) for more details.

**Table S9:** Age group comparison for behavioral and neural indicator variables

| Variable | nYA, nOA | *df* | *t* | *p* |
|---|---|---|---|---|
| Stay-standard | 36, 32 | 66 | 0.857 | 0.395 |
| Switch-to-standard | 36, 32 | 66 | 0.786 | 0.435 |
| Stay-oddball | 36, 32 | 66 | -0.273 | 0.785 |
| Switch-to-oddball | 36, 32 | 66 | -1.692 | 0.095 |
| Locus coeruleus integrity | 36, 32 | 66 | 0.418 | 0.677 |
| Pupil dilation | 36, 32 | 66 | -2.360 | 0.021 |
| Insula BOLD | 36, 32 | 66 | 0.797 | 0.429 |

*Note*: Younger adults (YA) were contrasted against older adults (OA). Thus, negative *t*-values indicate larger scores for older adults. nYA and nOA indicate participants per group entering the analyses. See Methods section (partial least squares analyses) for more details. Stay and switch variables indicate behavioral responses for trial categories of the conditioned oddball task. Uncorrected *p*-values are reported, with none of the statistics surviving correction for multiple comparisons.

#### Multimodal EEG–MRI analyses

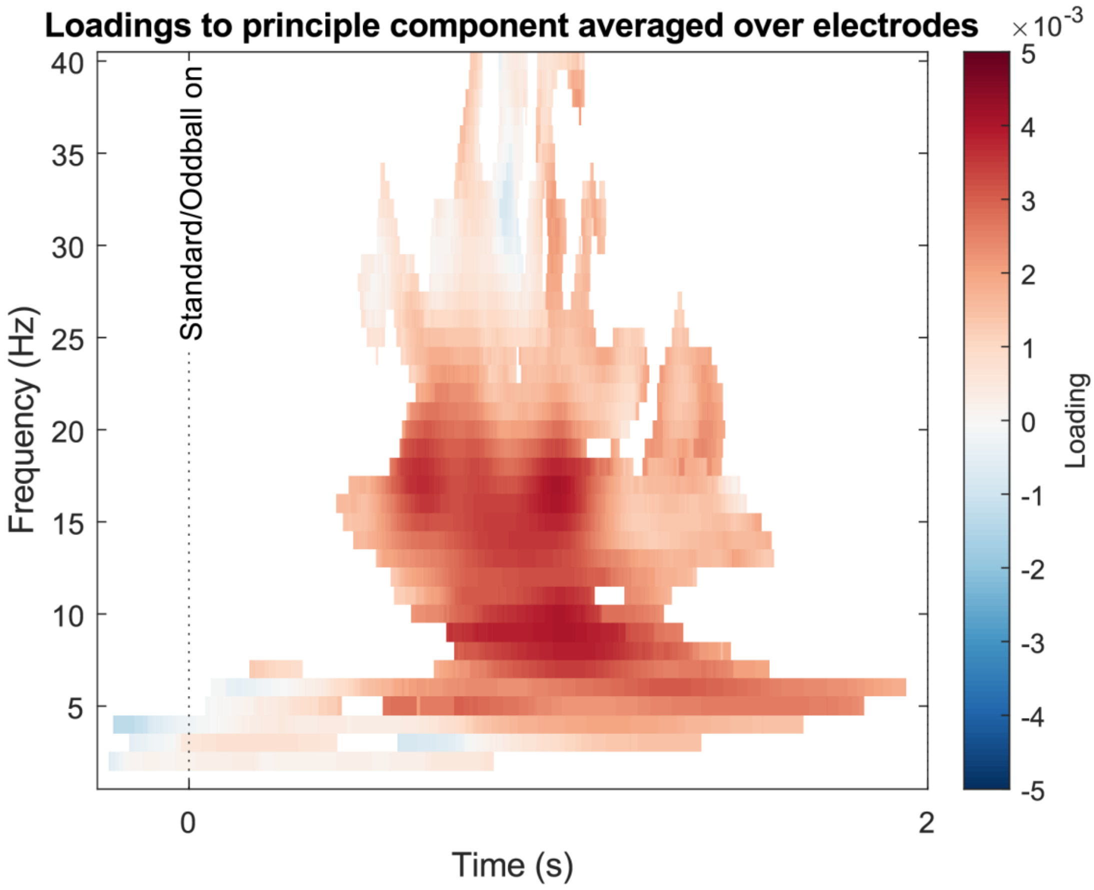

**Figure S11.** Sample-wise loadings to the principle component capturing individual differences in pupil-linked cortical excitability modulation strength. Highly similar loading patterns were obtained for the principle component analyses related to (1) *Combined EEG–MRI analyses* and (2) *Combined brain–behavior analyses including pupillometry and EEG* (see main text *Methods* section).

**Table S10:** Overlap of EEG–MRI Partial Least Squares Correlation-derived *positive* Bootstrap Ratios with canonical functional MRI networks

| Network name | Atlas | Dice coefficient | *p* |
|---|---|---|---|
| **Somato-motor** | **MG360J12** | 0.362 | 0.001 |
| **Action mode** | **MG360J12** | 0.323 | 0.006 |
| **Action mode** | **EG17** | 0.278 | 0.003 |
| **Dorsal attention** | **EG17** | 0.24 | 0.001 |
| Visual 2 | MG360J12 | 0.228 | 0.446 |
| Somato-motor A | TY17 | 0.223 | 0.048 |
| Fronto-parietal | EG17 | 0.212 | 0.231 |
| Fronto-parietal | MG360J12 | 0.209 | 0.738 |

*Note*: Functional networks are sorted in descending order of their overlap (Dice coefficient) with the EEG–MRI Partial Least Squares Correlation-derived loading pattern. Only networks with coefficients ≥ 0.2 are listed and only voxels with positive loadings are considered in the overlap estimation (>0). The following atlases are used: MG360J12 (Mattew Glasser2016 360-ROI with Ji2019 12 Cole-Anticevic networks), EG17 (Evan Gordon2017 17 networks), TY17 (Thomas Yeo 17 networks). Note that the Action-mode network has previously been termed Cingulo-opercular network and the Salience network has also been termed Ventral attention network.

**Table S11:** Overlap of EEG–MRI Partial Least Squares Correlation-derived *negative* Bootstrap Ratios with canonical functional MRI networks

| Network name | Atlas | Dice coefficient | *p* |
|---|---|---|---|
| **Default mode** | **MG360J12** | **0.499** | **0.001** |
| **Default mode** | **EG17** | **0.497** | **0.001** |
| **Default mode A** | **TY17** | **0.314** | **0.001** |
| **Default mode B** | TY17 | **0.286** | **0.001** |
| Fronto-parietal | MG360J12 | 0.24 | 0.341 |
| Visual 2 | MG360J12 | 0.214 | 0.483 |
| Lateral visual | EG17 | 0.208 | 0.235 |
| **Language** | **EG17** | **0.2** | **0.002** |

*Note*: Functional networks are sorted in descending order of their overlap (Dice coefficient) with the EEG–MRI Partial Least Squares Correlation-derived loading pattern. Only networks with coefficients ≥ 0.2 are listed and only voxels with negative loadings are considered in the overlap estimation (<0). The following atlases are used: MG360J12 (Mattew Glasser2016 360-ROI with Ji2019 12 Cole-Anticevic networks), EG17 (Evan Gordon2017 17 networks), TY17 (Thomas Yeo 17 networks). Note that the Action-mode network has previously been termed Cingulo-opercular network and the Salience network has also been termed Ventral attention network.

#### 3. References

Abraham, Alexandre, Fabian Pedregosa, Michael Eickenberg, Philippe Gervais, Andreas Mueller, Jean Kossaifi, Alexandre Gramfort, Bertrand Thirion, and Gael Varoquaux. 2014. "Machine Learning for Neuroimaging with Scikit-Learn." *Frontiers in Neuroinformatics* 8. https://doi.org/10.3389/fninf.2014.00014.

Avants, B.B., C.L. Epstein, M. Grossman, and J.C. Gee. 2008. "Symmetric Diffeomorphic Image Registration with Cross-Correlation: Evaluating Automated Labeling of Elderly and Neurodegenerative Brain." *Medical Image Analysis* 12 (1): 26–41. https://doi.org/10.1016/j.media.2007.06.004.

Behzadi, Yashar, Khaled Restom, Joy Liau, and Thomas T. Liu. 2007. "A Component Based Noise Correction Method (CompCor) for BOLD and Perfusion Based fMRI." *NeuroImage* 37 (1): 90–101. https://doi.org/10.1016/j.neuroimage.2007.04.042.

Cox, Robert W., and James S. Hyde. 1997. "Software Tools for Analysis and Visualization of fMRI Data." *NMR in Biomedicine* 10 (4-5): 171–78. https://doi.org/10.1002/(SICI)1099-1492(199706/08)10:4/5<171::AID-NBM453>3.0.CO;2-L.

Dale, Anders M., Bruce Fischl, and Martin I. Sereno. 1999. "Cortical Surface-Based Analysis: I. Segmentation and Surface Reconstruction." *NeuroImage* 9 (2): 179–94. https://doi.org/10.1006/nimg.1998.0395.

Esteban, Oscar, Ross Blair, Christopher J. Markiewicz, Shoshana L. Berleant, Craig Moodie, Feilong Ma, Ayse Ilkay Isik, et al. 2018. "FMRIPrep." *Software*. Zenodo. https://doi.org/10.5281/zenodo.852659.

Esteban, Oscar, Christopher Markiewicz, Ross W Blair, Craig Moodie, Ayse Ilkay Isik, Asier Erramuzpe Aliaga, James Kent, et al. 2018. "fMRIPrep: A Robust Preprocessing Pipeline for Functional MRI." *Nature Methods*. https://doi.org/10.1038/s41592-018-0235-4.

Evans, AC, AL Janke, DL Collins, and S Baillet. 2012. "Brain Templates and Atlases." *NeuroImage* 62 (2): 911–22. https://doi.org/10.1016/j.neuroimage.2012.01.024.

Fonov, VS, AC Evans, RC McKinstry, CR Almli, and DL Collins. 2009. "Unbiased Nonlinear Average Age-Appropriate Brain Templates from Birth to Adulthood." *NeuroImage* 47, Supplement 1: S102. https://doi.org/10.1016/S1053-8119(09)70884-5.

Gorgolewski, K., C. D. Burns, C. Madison, D. Clark, Y. O. Halchenko, M. L. Waskom, and S. Ghosh. 2011. "Nipype: A Flexible, Lightweight and Extensible Neuroimaging Data Processing Framework in Python." *Frontiers in Neuroinformatics* 5: 13. https://doi.org/10.3389/fninf.2011.00013.

Gorgolewski, Krzysztof J., Oscar Esteban, Christopher J. Markiewicz, Erik Ziegler, David Gage Ellis, Michael Philipp Notter, Dorota Jarecka, et al. 2018. "Nipype." *Software*. Zenodo. https://doi.org/10.5281/zenodo.596855.

Greve, Douglas N, and Bruce Fischl. 2009. "Accurate and Robust Brain Image Alignment Using Boundary-Based Registration." *NeuroImage* 48 (1): 63–72. https://doi.org/10.1016/j.neuroimage.2009.06.060.

Huntenburg, Julia M. 2014. "Evaluating Nonlinear Coregistration of BOLD EPI and T1w Images." Master's Thesis, Berlin: Freie Universität. http://hdl.handle.net/11858/00-001M-0000-002B-1CB5-A.

Jenkinson, Mark, Peter Bannister, Michael Brady, and Stephen Smith. 2002. "Improved Optimization for the Robust and Accurate Linear Registration and Motion Correction of Brain Images." *NeuroImage* 17 (2): 825–41. https://doi.org/10.1006/nimg.2002.1132.

Klein, Arno, Satrajit S. Ghosh, Forrest S. Bao, Joachim Giard, Yrjö Häme, Eliezer Stavsky, Noah Lee, et al. 2017. "Mindboggling Morphometry of Human Brains." *PLOS Computational Biology* 13 (2): e1005350. https://doi.org/10.1371/journal.pcbi.1005350.

Lanczos, C. 1964. "Evaluation of Noisy Data." *Journal of the Society for Industrial and Applied Mathematics Series B Numerical Analysis* 1 (1): 76–85. https://doi.org/10.1137/0701007.

Power, Jonathan D., Anish Mitra, Timothy O. Laumann, Abraham Z. Snyder, Bradley L. Schlaggar, and Steven E. Petersen. 2014. "Methods to Detect, Characterize, and Remove Motion Artifact in Resting State fMRI." *NeuroImage* 84 (Supplement C): 320–41. https://doi.org/10.1016/j.neuroimage.2013.08.048.

Pruim, Raimon H. R., Maarten Mennes, Daan van Rooij, Alberto Llera, Jan K. Buitelaar, and Christian F. Beckmann. 2015. "ICA-AROMA: A Robust ICA-Based Strategy for Removing Motion Artifacts from fMRI Data." *NeuroImage* 112 (Supplement C): 267–77. https://doi.org/10.1016/j.neuroimage.2015.02.064.

Satterthwaite, Theodore D., Mark A. Elliott, Raphael T. Gerraty, Kosha Ruparel, James Loughead, Monica E. Calkins, Simon B. Eickhoff, et al. 2013. "An improved framework for confound regression and filtering for control of motion artifact in the preprocessing of resting-state functional connectivity data." *NeuroImage* 64 (1): 240–56. https://doi.org/10.1016/j.neuroimage.2012.08.052.

Treiber, Jeffrey Mark, Nathan S. White, Tyler Christian Steed, Hauke Bartsch, Dominic Holland, Nikdokht Farid, Carrie R. McDonald, Bob S. Carter, Anders Martin Dale, and Clark C. Chen. 2016. "Characterization and Correction of Geometric Distortions in 814 Diffusion Weighted Images." *PLOS ONE* 11 (3): e0152472. https://doi.org/10.1371/journal.pone.0152472.

Tustison, N. J., B. B. Avants, P. A. Cook, Y. Zheng, A. Egan, P. A. Yushkevich, and J. C. Gee. 2010. "N4ITK: Improved N3 Bias Correction." *IEEE Transactions on Medical Imaging* 29 (6): 1310–20. https://doi.org/10.1109/TMI.2010.2046908.

Wang, Sijia, Daniel J. Peterson, J. C. Gatenby, Wenbin Li, Thomas J. Grabowski, and Tara M. Madhyastha. 2017. "Evaluation of Field Map and Nonlinear Registration Methods for Correction of Susceptibility Artifacts in Diffusion MRI." *Frontiers in Neuroinformatics* 11. https://doi.org/10.3389/fninf.2017.00017.

Zhang, Y., M. Brady, and S. Smith. 2001. "Segmentation of Brain MR Images Through a Hidden Markov Random Field Model and the Expectation-Maximization Algorithm." *IEEE Transactions on Medical Imaging* 20 (1): 45–57. https://doi.org/10.1109/42.906424.

Zeidman, P., Jafarian, A., Corbin, N., Seghier, M. L., Razi, A., Price, C. J., & Friston, K. J. (2019). "A guide to group effective connectivity analysis, part 1: First level analysis with DCM for fMRI." *NeuroImage*, 200, 174–190. https://doi.org/10.1016/J.NEUROIMAGE.2019.06.031
